# Single-cell spatial transcriptomics of fixed, paraffin-embedded biopsies reveals colitis-associated cell networks

**DOI:** 10.1101/2024.11.11.623014

**Authors:** Elvira Mennillo, Madison L. Lotstein, Gyehyun Lee, Vrinda Johri, Christina Ekstrand, Jessica Tsui, Julian Hou, Donna E. Leet, Jun Yan He, Uma Mahadevan, Walter Eckalbar, David Y. Oh, Gabriela K. Fragiadakis, Michael G. Kattah, Alexis J. Combes

## Abstract

**Background & Aims:** Imaging-based, single-cell spatial transcriptomics (iSCST) using formalin-fixed, paraffin-embedded (FFPE) tissue could transform translational research by retaining all tissue cell subsets and spatial locations while enabling the analysis of archived specimens. We aimed to develop a robust framework for applying iSCST to archived clinical FFPE mucosal biopsies from patients with inflammatory bowel disease (IBD).

**Methods:** We performed a comprehensive benchmarking comparison of three iSCST platforms capable of analyzing FFPE specimens. We analyzed FFPE mucosal biopsies (n=57) up to 5 years old from non-IBD controls (HC; n=9) and patients with ulcerative colitis (UC;n=11). After platform-specific cell segmentation, we applied a uniform data processing pipeline to all datasets, including transcript detection, cell annotation, differential gene expression, and neighborhood enrichment. Transcriptomic signatures identified with iSCST were validated using external, publicly available bulk transcriptomic datasets.

**Results:** A custom 290-plex Xenium gene panel exhibited the highest sensitivity and specificity for transcript detection, enabling precise identification and quantification of diverse cell subsets and differentially expressed genes across cell types and disease states. We mapped transcriptionally distinct fibroblast subsets to discrete spatial locations and identified inflammation-associated fibroblasts (IAFs) and monocytes as a colitis-associated cellular neighborhood. We also identified signatures associated with Vedolizumab (VDZ) responsiveness. VDZ non-responders were characterized by an IAF-monocyte transcriptional signature, while responders exhibited enrichment of epithelial gene sets.

**Conclusions:** Our optimized iSCST framework for archived FFPE biopsies provides unique advantages for assessing the role of colitis-associated cellular networks in routinely collected clinical samples. FFPE-based biomarkers could integrate with existing clinical workflows and potentially aid in risk-stratifying patients.

## INTRODUCTION

Single-cell multi-omics studies in inflammatory bowel disease (IBD) have rapidly accelerated our understanding of intestinal inflammation, highlighting critical roles for multiple cell subsets across stromal, epithelial, and immune compartments^1–8^. While powerful, traditional single-cell multi-omics analyses of prospectively collected and digested colon biopsies suffer from several important limitations. These include cellular dropout or under-representation of epithelial and granulocyte subsets, loss of spatial information, and prolonged patient recruitment^9,10^. As an example, neutrophils contribute to acute colitis and treatment non-response, but they are particularly difficult to capture with many single-cell multi-omics approaches, and their proximity and interactions with inflammatory fibroblasts are lost in disaggregated tissue^8,10,11^. Analyzing cellular networks such as these before and after treatment in responders and non-responders across multiple therapies from prospectively collected samples requires prolonged recruitment. Spatial transcriptomics of formalin-fixed, paraffin-embedded (FFPE) tissue with subcellular resolution directly addresses these shortcomings by comprehensively capturing cell subsets and transcriptional states in mucosal biopsies, retaining spatial relationships, and accelerating recruitment for retrospective, longitudinal, case-control analyses of archived clinical specimens collected as part of standard of care.

Spatial transcriptomics technologies hold great promise for identifying tissue biomarkers and augmenting our understanding of disease and treatment response. These technologies are still emerging, and a growing body of literature leverages these approaches to study mouse models^12^ or patient samples^10,13^, but few studies have benchmarked different platforms for the optimal analysis of archived FFPE clinical specimens, and those that have were primarily focused on tumor tissues^14–17^. Hence, it is unclear whether prior iSCST comparisons extrapolate to archived clinical specimens from inflammatory disorders such as IBD. We performed a comparison of three commercially available imaging-based single-cell spatial transcriptomics (iSCST) platforms capable of analyzing FFPE specimens with subcellular resolution from tissue sections. These platforms were used to analyze FFPE colon mucosal biopsies archived for up to 5 years from non-IBD controls and patients with UC. Our primary comparison examined specific and non-specific transcript detection, cell-subset identification, and differential gene expression between healthy and inflamed tissue. We also examined cell segmentation and the impact of sample age on assay performance. Using our custom IBD panel, we identified transcriptionally and spatially distinct fibroblast subsets in health and disease, and we defined an inflammation-associated fibroblast (IAF)-monocyte cellular network that increased in abundance and spatial proximity in patients with active UC. We present an optimized iSCST pipeline for analyzing archived mucosal biopsies from patients with inflammatory intestinal disorders, offering unique advantages for evaluating colitis-associated cellular networks in routinely collected clinical samples.

## RESULTS

### Xenium Outperforms Competing iSCST Platforms in Sensitivity and Detection of Colon-Associated Cell Types

To develop an iSCST pipeline for studying archived FFPE specimens, here focused on mucosal biopsies from IBD patients, we evaluated three commercially available imaging-based spatial transcriptomics platforms: Xenium (10X Genomics), CosMx (NanoString), and MERSCOPE (Vizgen). The CosMx dataset was previously generated using a 1,000 gene predesigned CosMx Human Universal Cell Characterization RNA Panel on an FFPE tissue microarray (TMA) containing colon biopsies from UC patients and patients without IBD, referred to as healthy controls (HC), and was re-analyzed for this study (**Figure 1A, Supplementary Table 1**)^9^. The Xenium and MERSCOPE spatial platforms were used to test the same TMA that was used to generate the CosMx data. The custom Xenium panel contained 290 genes, and the custom MERSCOPE panel contained 280 genes (**Supplementary Table 2)**. Both panels were designed to detect colon-associated mucosal and submucosal cell subsets and differentially expressed genes (DEGs) identified by single-cell RNA sequencing (scRNA-seq) of colon biopsies from UC and HC patients^9^. Due to difficulties with tissue clearing, MERSCOPE could not detect most of the genes in the panel (**Supplementary Figure 1A-B**). Even though some genes were detected as shown in the TMA scanned area (**Supplementary Figure 1C**), they were low quality and only 0.17% of cells passed quality control (QC) (**Supplementary Figure 1D**). Thus, MERSCOPE was excluded from the remainder of the analysis. The Xenium and CosMx panels shared 159 overlapping genes and were compared for transcript and gene detection, cell annotation, and differential gene expression (**Supplementary Table 2)**.

**Fig. 1.**
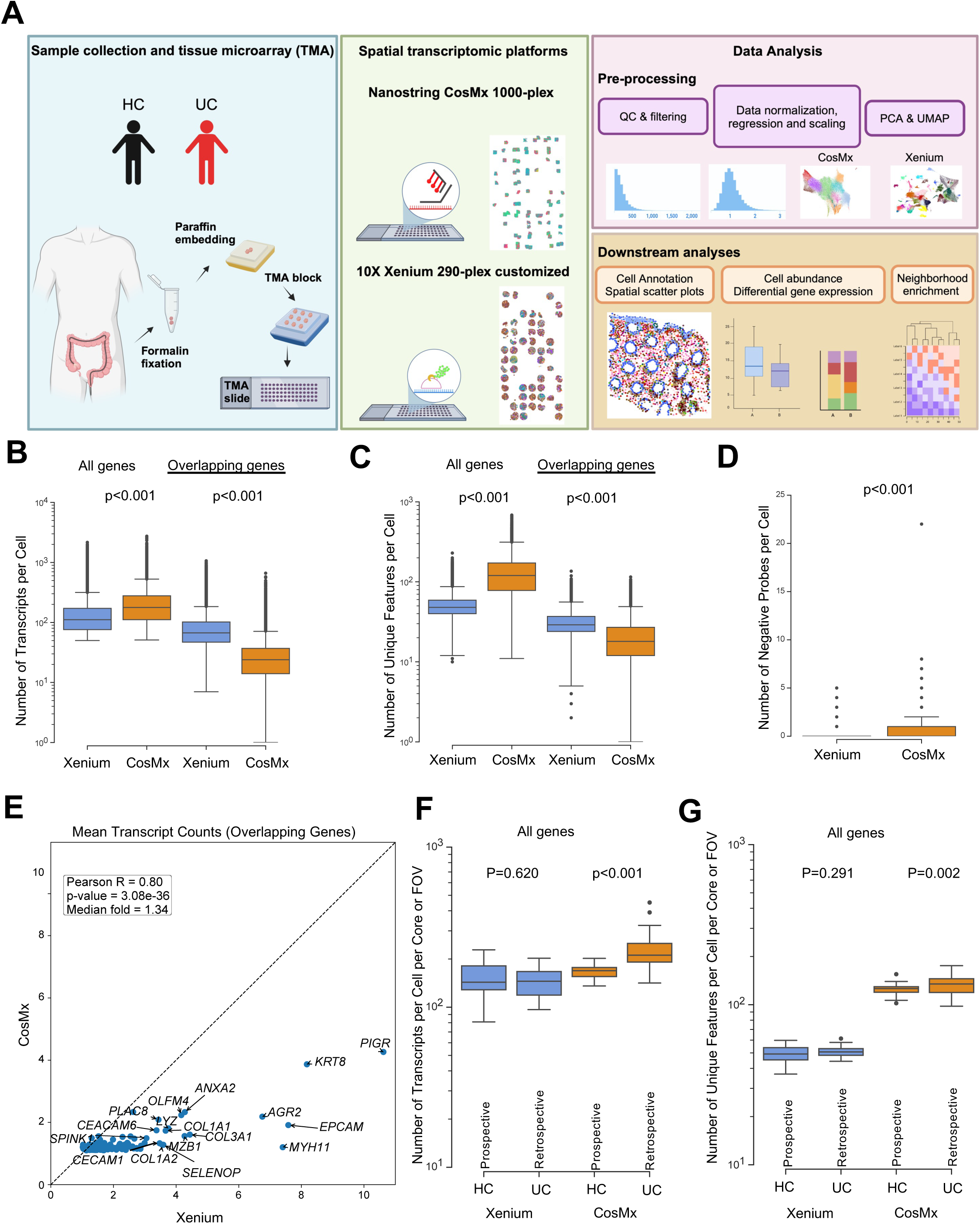
Schematic of study design and technical performance comparison between Xenium and CosMx platforms. (**A**) Schematic of study design. Created with BioRender.com (**B,C**) Number of (**B**) transcripts and (**C**) unique features detected per cell within the Xenium and CosMx datasets, calculated using the complete gene panel for each platform (Xenium, 290 genes; CosMx, 1,000 genes; left) and limited to the 159 overlapping genes across both panels (right). (**D**) Number of negative probes detected per cell within the Xenium and CosMx (Xenium mean=0.03 and CosMx mean=0.37). (**E**) Correlation between average transcript counts in Xenium and CosMx for the 159 overlapping genes. (**F,G**) Number of (**F**) transcripts and (**G**) unique features detected per cell per core or FOV within the Xenium and CosMx datasets split by prospectively collected HC and retrospectively collected UC. For panels **B, C, D, F** and **G** box and whisker plots, the band indicates the median, the box indicates the first and third quartiles, and the whiskers indicate minimum and maximum value within the upper/lower fence (upper fence=Q3+1.5xIQR and lower fence= Q1-1.5xIQR), only outlier points are shown. Mann-Whitney, two-tailed tests, p-values are indicated.

We began by using the default cell segmentation settings for each platform to identify single cells. For the Xenium platform, we employed the integrated default nuclear expansion method, extending nuclear boundaries by 5 μm or until another cell boundary is encountered. In contrast, the CosMx platform combines nuclear and cell membrane staining for cell identification. Although this led to slight differences in the distribution of nuclear versus cytoplasmic transcripts between platforms (**Supplementary Figure 2A**), the median cell area was similar (**Supplementary Figure 2B**). To further investigate outlier cells with larger areas observed in the Xenium platform, we examined cell boundaries using the Xenium Explorer visualization tool from 10X Genomics. Some cells displayed expanded boundaries, but the distortion affected primarily cells adjacent to luminal space (**Supplementary Figure 2C**). Overall, the nuclear expansion cell segmentation inflated the cell area of a relatively small percentage of cells.

We then evaluated sensitivity metrics across the full gene panel for both the Xenium and CosMx platforms. The larger CosMx panel yielded a higher number of transcripts and unique features per cell (**Figure 1B-C**). However, when the analysis was restricted to the 159 overlapping genes between the two panels, Xenium detected significantly more transcripts and unique features per cell compared to CosMx (**Figure 1B-C**). Importantly, Xenium also exhibited ten times lower detection of negative probes per cell (0.03 vs 0.37, p<0.001) (**Figure 1D**). Xenium demonstrated significantly higher transcript counts for the overlapping genes (**Figure 1E**). The higher signal per gene with reduced non-specific detection of negative probes yielded a markedly higher signal over background for Xenium compared to CosMx (**Supplementary Table 3**). Upon examining dataset-specific genes unique to either the Xenium or CosMx panels, we found that most genes exhibited similar expression levels, typically ranging from 1 to 5 transcripts per cell. However, immunoglobulin genes expressed by B and plasma cells (*IGHG2, IGKC, IGHG1, IGHA1*) displayed higher expression levels (**Supplementary Figure 2D**). These highly expressed immunoglobulin genes could not be included in the Xenium panel due to their abundance and issues with optical crowding, pointing to one limitation of the signal amplification with Xenium. Notably, the age of the sample block had minimal impact on performance metrics for both methods, showing no meaningful differences in the median number of transcripts or genes detected per cell per core or FOV across 2-, 3-, 4-, and 5-year-old sample blocks (**Supplementary Figure 2E-F**). The HC samples were prospectively collected and processed in the laboratory, while the UC samples were retrospectively retrieved years after processing and storage through the usual standard of care. Despite this, the retrospectively retrieved UC samples exhibited equivalent or better transcript and feature detection compared to prospectively collected HC samples (**Figure 1F-G)**.

A key advantage of iSCST is its ability to spatially locate transcriptomic cell states that were previously identified by unsupervised clustering of single-cell RNA sequencing (scRNA-seq) from dissociated tissue. We assessed how each platform’s specificity and sensitivity affected unsupervised cell type identification in FFPE colon biopsies. Using the same data processing pipeline, we generated uniform manifold approximation and projection (UMAP), clustering, and annotation, and generated heatmaps of landmark genes and correlation matrices for Xenium (**Fig 2A-D**) and CosMx (**Fig 2E-H**). At both the coarse (**Figure 2A, 2E**) and fine level (**Figure 2B, 2F**), comparing this CosMx re-analysis and prior analysis^9^, Xenium enabled the identification of a broader range of colon-associated cell types, including various fine subsets of epithelial cells, fibroblasts, endothelial cells, and fragile innate immune cells like neutrophils and mast cells. Some subsets were difficult to confidently assign, such as intraepithelial lymphocytes (IELs), which is likely due in part to cell segmentation challenges. Overall, unsupervised clustering of single-cell gene expression profiles from Xenium produced more distinct gene expression signatures per cluster, yielding clearer cell cluster separation, as shown in heatmaps (**Figure 2C, 2G**) and correlation plots (**Figure 2D, 2H**). In contrast, the same data processing pipeline of the CosMx data from the same TMA resulted in >20% of cells being labeled as “Unassigned” due to indistinct transcriptomic profiles (**Figure 2I**). There was also a small percentage of cells labeled “ncount_hi” characterized by the high expression of B and plasma cell-related genes. Although finer cell annotation is possible with both the Xenium and CosMx datasets, here we applied comparable pipelines to provide a more direct comparison of subset identification. We then assessed the mutually exclusive co-expression rate (MECR) values for each platform, which provides a metric for lineage-specific genes that are unique to prominent cell types (**Supplementary Figure 3**). As a gold standard, we compared MECR values to previously published scRNA-seq data from HC and UC patient biopsies^9^. MECR values were generally similar for the two iSCST platforms, and neither was as specific as scRNA-seq, but the higher transcript counts for unique landmark genes enabled finer cell annotation with Xenium (**Supplementary Figure 3**, **Figure 2B**). Overall, the improved performance of Xenium for cell identification in these colon FFPE samples is due to multiple factors, including higher cell number, a custom panel optimized for colon biopsies, and higher sensitivity and specificity for landmark gene expression. Then, to address reproducibility with the Xenium platform, we performed a replicate run of separate sections from the same TMA. We observed high reproducibility for gene expression levels (**Supplementary Figure 4A**) and the samples demonstrated a balanced representation of cell types across replicates (**Supplementary Figure 4B**). Doubling the number of cells in our dataset enhanced the accuracy of our detailed fine annotations (**Supplementary Figure 4C, Supplementary Figure 5A-B**). Taken together, our comprehensive comparison of three iSCST platforms for FFPE specimens indicates that the Xenium platform exhibits superior transcript and gene sensitivity, enables the detection of a greater range of colon-associated cell types and transcriptional states, exhibits high reproducibility across replicates, and is robust to tissue age.

**Fig. 2.**
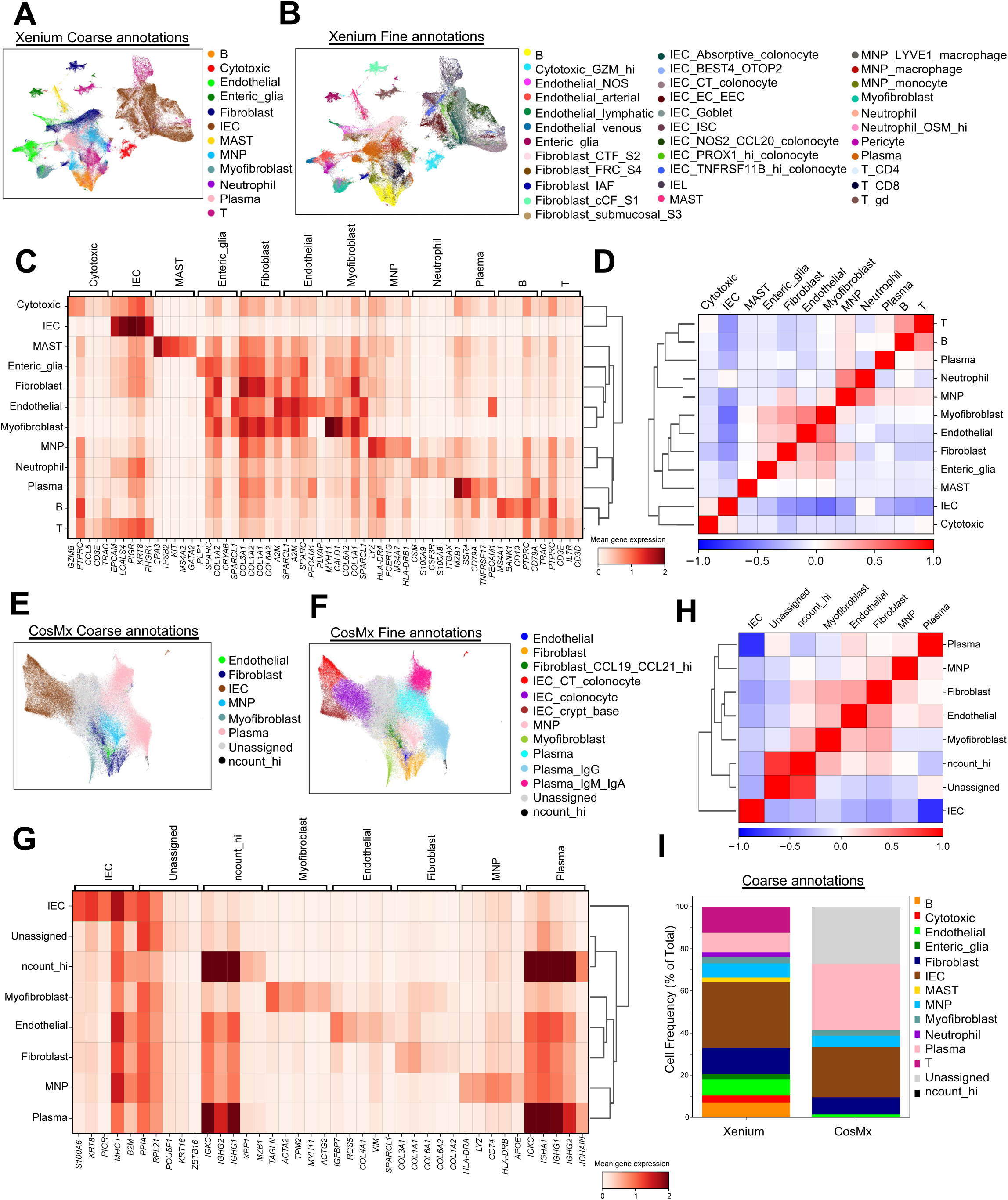
Cell type recovery across spatial transcriptomics platforms. (**A,B**) UMAP visualization of Xenium dataset (313,940 cells), colored by coarse (**A**) and fine (**B**) annotations. (**C**) Heatmap displaying gene expression of the top 5 landmark genes for each coarse annotation cell type within the Xenium dataset. (**D**) Correlation matrix displaying the correlation between coarse annotation cell types identified for Xenium. (**E,F**) UMAP visualization of CosMx dataset (126,368 cells), colored by coarse (**E**) and fine (**F**) annotations. (**G**) Heatmap displaying gene expression of the top 5 landmark genes for each coarse annotation cell type within the CosMx dataset. (**H**) Correlation matrix displaying the correlation between coarse annotation cell types identified for CosMx. (**I**) Stacked bar plots for coarse annotation displaying cell frequency (percent of total) for Xenium and CosMx.

### A custom 290-plex iSCST Xenium panel differentiates cases and controls in UC FFPE colon biopsies, revealing spatial transcriptomic profiles of B Cells, epithelial cells, and mononuclear phagocytes

Next, we aimed to extend our comparison of the CosMx and Xenium iSCST platforms beyond QC and cell identification metrics to evaluate their ability to identify transcriptomic signatures associated with UC pathology. Publicly available bulk transcriptomic data served as the comparator for this analysis, rather than scRNA-seq, because bulk RNA isolation methods do not involve digestion and, therefore, do not suffer from cellular or transcriptional dropout like scRNA-seq studies. We generated pseudobulk iSCST transcriptomic profiles by aggregating the transcript counts for all cells for each individual patient. We then performed a DEG analysis comparing UC biopsies before treatment with VDZ (UC PRE) to HC using the full gene panels for both platforms. Using a cutoff of >0.4 Log2FC and q-value <0.1, the transcriptomic profile obtained from the Xenium platform identified a broad range of upregulated genes in both UC and HC groups, with 32 genes significantly upregulated in UC PRE and 60 in HC (**Figure 3A**). The UC-associated DEGs included genes well-known for their association with inflammatory infiltrates, specifically neutrophils and monocytes (S100A8/S100A9) as well as lymphocytes (CD19, LTB, SELL, MS4A1), with cell identities confirmed in our Xenium iSCST dataset (**Supplementary Figure 5C**). Conversely, HC-associated genes included mostly genes associated with intact intestinal epithelial cells (IECs), such as *c15orf48, EPCAM*, *AQP8*, *FABP1, FABP2, PHGR1* (**Supplementary Figure 5C**). In contrast, data from the CosMx platform on the same TMA identified 57 genes upregulated in UC PRE and only one gene significantly upregulated in HC (**Figure 3B**). Unsupervised hierarchical clustering of a subset of the DEGs showed unique expression patterns that readily distinguished UC PRE patients and HC in the Xenium dataset (**Figure 3C**) but not CosMx (**Figure 3D**). CosMx identified a slightly higher number of DEGs in the UC PRE samples compared to Xenium, but only identified a single gene associated with HC (**Figure 3E**). It is important to acknowledge that testing more genes (1000 vs 290) leads to a statistical penalty for CosMx due to FDR correction. We then focused our comparison on the overlapping genes across platforms. This revealed a poor correlation between the differentially expressed genes identified by the two platforms (R = 0.12), with only *COL1A1*, *MZB1*, and *TIMP1* being significant in both datasets (**Figure 3F**). Overall, genes identified by Xenium exhibited a higher fold change compared to those from CosMx (**Figure 3F**). To further validate our Xenium gene signatures identified for HC and UC PRE, we performed gene set enrichment analysis (GSEA) on an external bulk transcriptomic dataset^18^, confirming significant Normalized Enrichment Scores (NES) in HC and UC PRE, and nearly flawless validation of the UC PRE-associated signature (**Figure 3G**). CosMx showed a slightly lower NES compared to Xenium for the UC PRE signature (−2.49 vs −2.66; **Figure 3G** and **Supplementary Figure 6A)**. To further adjudicate the conflicting DEGs between the Xenium and CosMx datasets (**Figure 3F**), we then determined whether the individual DEGs from the two iSCST methods exhibited concordance or discordance with up-regulation in the UC PRE external bulk transcriptomic dataset. Although both methods revealed significant concordance, CosMx demonstrated more discordance with bulk transcriptomic data, as 13 DEGs in the CosMx UC PRE signature were instead associated with HC in bulk transcriptomic data (**Supplementary Figure 6B**). These discordant CosMx genes included *KRT20* and *LGALS3*, which were enriched in HC in the external dataset and are expressed by intact IECs (**Supplementary Table 5**).

**Fig. 3.**
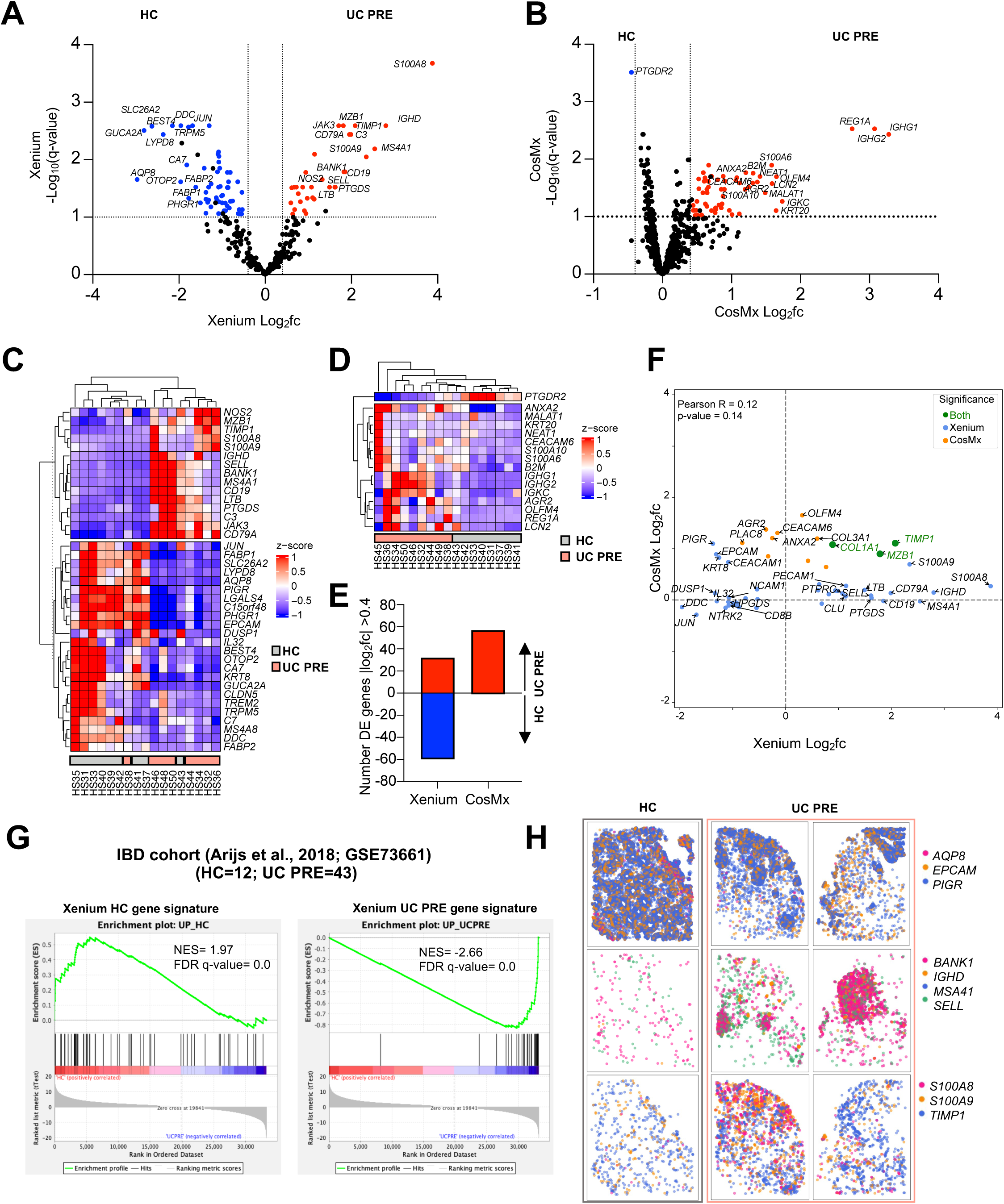
Pseudobulk DE gene analysis comparing UC PRE versus HC in both platforms and gene set enrichment analysis (GSEA) of an external, publicly available, bulk transcriptomic dataset using Xenium transcriptomic signatures. (**A**) Volcano plot of pseudobulk DE genes identified by DESeq2 with log_2_fc >0.4 or <-0.4 and q < 0.1 in UC PRE vs HC for Xenium dataset. (**B**) Volcano plot of pseudobulk DE genes identified by DESeq2 with log_2_fc >0.4 or <-0.4 and q < 0.1 in UC PRE vs HC for CosMx dataset. (**C**) Heatmap of expression z-scores for the indicated genes in UC PRE (Up/Down) relative to HC for Xenium dataset. (**D**) Heatmap of expression z-scores for the indicated genes in UC PRE (Up/Down) relative to HC for CosMx dataset. (**E**) Number of pseudobulk DE genes in the indicated platform with log2fc > 0.4 or <-0.4 in UC PRE relative to HC identified by DESeq2 analysis. (**F**) Volcano plot of significant overlapping genes identified by DESeq2 with log_2_fc >0.4 or <-0.4 for Xenium and CosMx; genes are color coded (green, significant in both panels; blue, significant in Xenium panel only; orange, significant in CosMx panel only). (**G**) GSEA of Xenium HC and UC PRE spatial gene signatures in an external cohort of patients and relative Normalized Enrichment Scores (NES). (**H**) Representative spatial transcript scatter plots highlighting a subset of genes relatively increased in HC and UC PRE in Xenium dataset. For panels **B** and **D**, some genes are off-scale for visualization purposes, z-score set from −1 to 1.

Finally, we leveraged the spatial information associated with each transcript to create spatial transcript scatter plots for the mucosal biopsies for subsets of DEGs identified by Xenium for HC and UC PRE patients. Transcript-level spatial scatter plots for IEC-specific genes (*AQP8, EPCAM*, and *PIGR*) were more abundant in HC tissue, while B cell-specific genes (*BANK1, IGHD, MS4A1*, and *SELL*) and MNP and neutrophil-specific genes (*S100A8, S100A9*, and *TIMP1*) were linked to disease (**Figure 3H**). Overall, the custom Xenium 290-plex gene panel identified more DEGs between HC and UC patients, with higher log fold changes and greater concordance with external validation bulk transcriptomic data. This strong differentiation between cases and controls in UC FFPE colon biopsies is marked by a shift in transcriptomic profiles, from epithelial cells predominating in HC to B cells, neutrophils and MNPs in UC.

### Optimized iSCST pipeline identifies spatially and transcriptionally distinct fibroblast populations in FFPE colon biopsies

Previous work by our group^9^ and others^1,19^ has highlighted the importance of distinct fibroblast subsets, characterized by unique transcriptional states, in colon tissue under steady-state conditions, as well as in UC pathology and treatment response. While the spatial location of those transcriptomic subsets has been recently described in IBD mouse models^12^, equivalent detailed spatial characterization in clinical biopsies has been challenging and is still lacking. By leveraging the high sensitivity of the Xenium iSCST platform, unsupervised clustering from our custom 290-plex gene panel identified five transcriptionally distinct fibroblast subsets (S1, S2, S3, S4 and IAF; **Figure 2B and Supplementary Figure 5A**) that were previously characterized in scRNA-seq of human colonic stromal cells^1,9,19^. In agreement with prior scRNA-seq studies, Xenium iSCST identified transcriptionally distinct S1 (*CCL8*, *ADAMDEC1, APOE, FABP5)*, S2 (*F3, SOX6, PDGFRA, WNT5A, POSTN, CXCL14*), S3 (*GSN, OGN, CCDC80, C7*), S4 (*C3, TNFSF13B, IRF8, PTGDS)*, and IAF (*IL1R1, TIMP1, CD44, IL13RA2, MMP1, MMP3, OSMR, NFKBIA, TNFAIP3*) subsets (**Figure 4A**). These subsets display a broad range of functions and identities, and some have previously been linked to specific locations in the colon^20^. S1 fibroblasts have been linked to *PDGFRA^lo^*colonic crypt fibroblasts (cCF) at the bottom of the colon crypts, S2 fibroblasts have been described to line the colonic crypt, and share features with *PDGFRA^hi^* crypt top fibroblasts (CTFs) that are also high in *SOX6* and *WNT5A*^1,9,19,20^. While IAF are disease-associated, S1-4 subsets were present at steady state. We spatially mapped the transcriptomic states present in colon tissue at steady state (**Figure 4B**). Spatial scatter plots showed that the S1-cCF fibroblasts were somewhat dispersed but were biased more toward the crypt base, while S2-CTF fibroblasts lined the colonic crypt as previously described (**Figure 4B**). S3 fibroblasts localize to the submucosal (**Figure 4B**), while S4 fibroblasts express fibroblastic reticular cell (FRC)-associated genes, including *TNFSF13B* or B-cell activating factor (BAFF) and colocalize with B cells in gut-associated lymphoid tissue (GALT) aggregates (**Figure 4B**). Together, this study provides a comprehensive spatial map of transcriptomic subsets of colonic fibroblasts at steady state in archived clinical FFPE mucosal biopsies using iSCST.

**Fig. 4.**
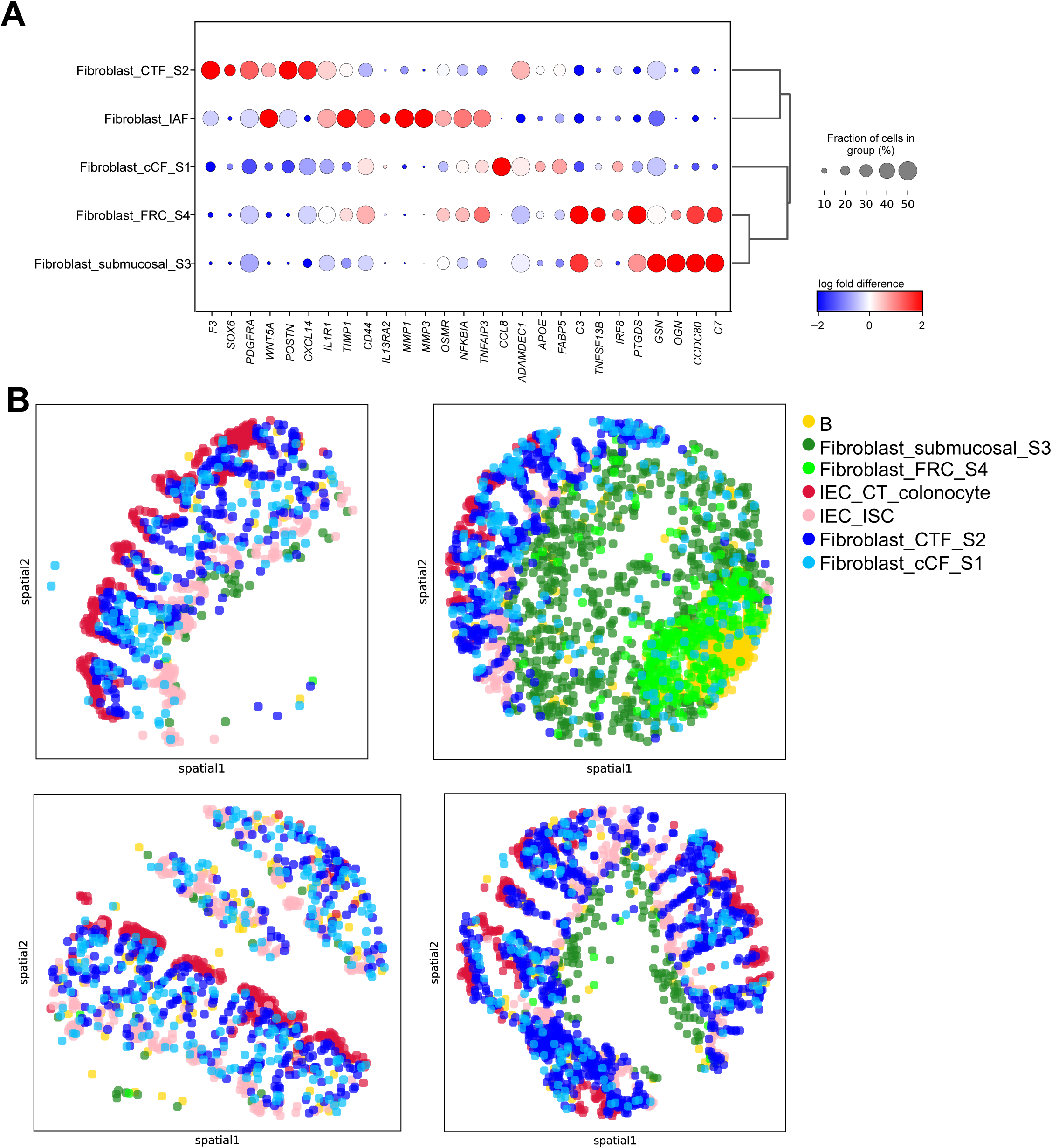
Xenium enabled identification and spatial localization of distinct fibroblast subsets in colon mucosal biopsies and identified increased myeloid and stromal cell subsets in UC. (A) Dot plot representation of landmark genes for the indicated subsets. (B) Transcriptionally distinct fibroblast subsets identified by relative spatial localization in colon tissue from representative cores for the indicated cell subsets.

### Inflammation-associated fibroblast–monocyte network and B cell abundance in GALT are associated with UC

Our spatially resolved Xenium iSCST dataset accurately identified 34 distinct transcriptomic cell states, including fragile populations such as neutrophils which are typically lysed during cryopreservation and thawing or are excluded by encapsulation in single-cell sequencing workflows^9^. This provided us with a unique opportunity to study cell composition, gene expression changes, and spatial location in undissociated colon tissue between HC and UC cases without cellular dropout. First, by comparing cell abundances between HC and UC PRE, we revealed 19 cell subsets that differed significantly after FDR correction (**Figure 5A and Supplementary Figure 7**). We observed significant variations associated with disease, specifically in monocyte, neutrophil, IAF, and S4 fibroblast abundance (**Figure 5A**). Furthermore, there was a significant increase across all endothelial subsets, pericytes, B and plasma cells, and both CD4 and CD8 T lymphocytes in UC PRE, with a reduction in IEC subsets (**Supplementary Figure 7**). iSCST with Xenium identified transcriptionally and spatially distinct colonic mucosal cell subsets in homeostasis and disease, providing a more comprehensive framework for interrogating stromal and immune subsets.

**Fig. 5.**
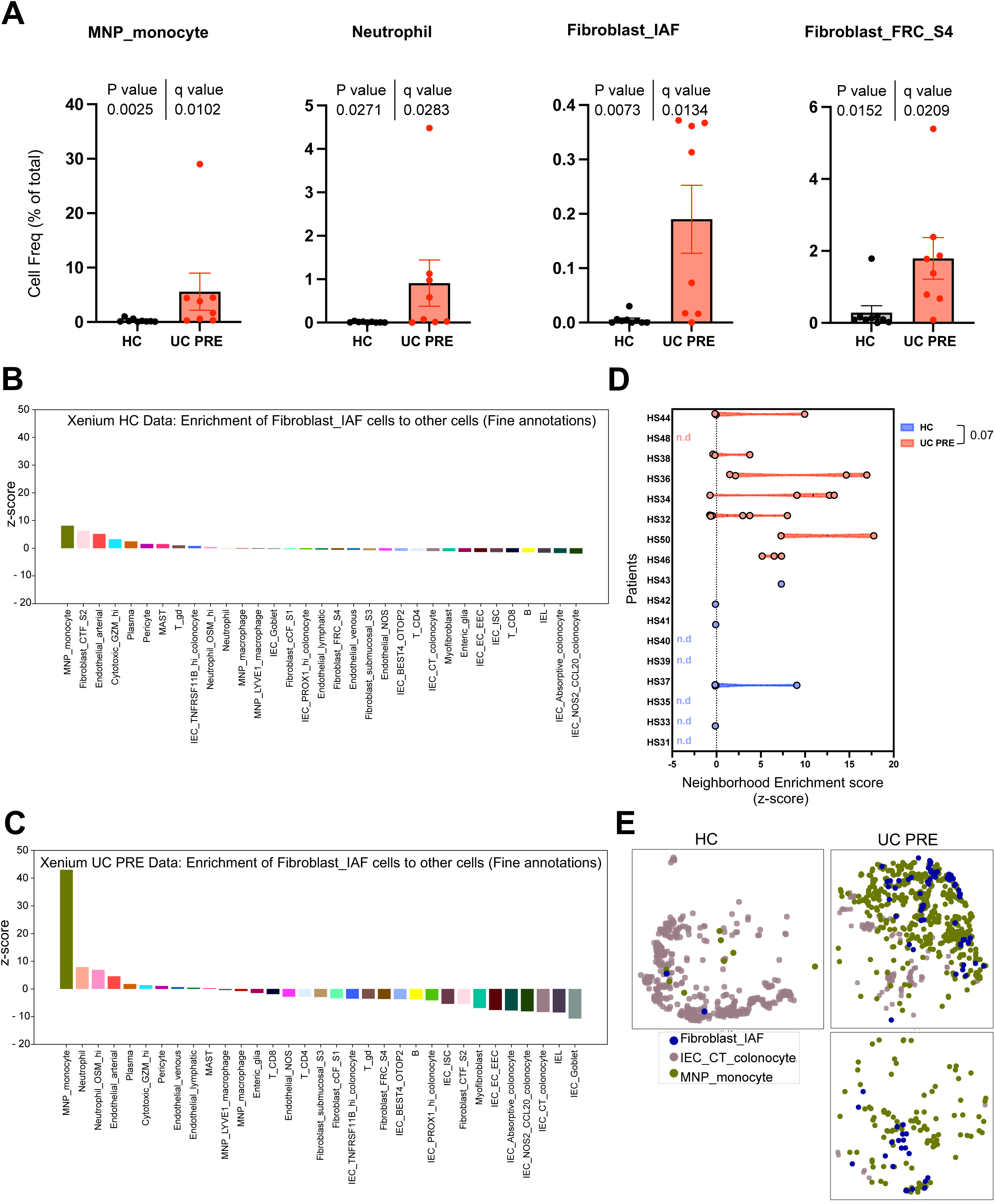
Neighborhood enrichment analysis of Xenium dataset reveals higher proximity of IAFs and monocytes in UC PRE biopsies. (**A,B**) Heatmaps displaying neighborhood enrichment z-scores for fine annotation cell pairs within (**A**) HC and (**B**) UC PRE biopsies. Spatial enrichment of IAFs cells to all other cell types in (**C**) HC and (**D**) UC PRE biopsies. (**E**) Violin plots comparing the spatial enrichment of IAFs and monocytes by patient, each dot represents a core; nd, not defined. (**F**) Spatial scatter plot of representative cores highlighting IAF, crypt top (CT) colonocytes, and monocytes in HC and UC PRE biopsies. For panel **E**, Mann-Whitney test, p-value is indicated. For panels **A** and **B**, several values exceed the scale for visualization purposes and are denoted by a white asterisk.

We then leveraged the spatial aspect of the dataset to explore the proximity and interactions between our annotated cell types, comparing HC and UC PRE. Spatial neighborhood graphs were constructed for the HC and UC PRE data subsets using the fine annotation cell type categories aggregating all patients. Using these spatial neighbor graphs, we calculated neighborhood enrichment z-scores for each cell type pair (**Supplementary Figure 8**). Comparing cellular neighborhoods between HC and UC PRE, we identified several notable changes for the monocytes, including neighborhood enrichment with IAFs, neutrophils, and B cells (**Supplementary Figure 8B**). Next, we focused on the neighborhood enrichment z-scores between IAFs and all other cell types in HC (**Figure 5B**) and UC PRE (**Figure 5C**). Monocytes exhibited the highest interaction scores in both HC and UC PRE, with a collectively higher value in UC PRE. The IAF-monocyte enrichment was followed by IAF-neutrophil enrichment, which has been described in recent single-cell analyses of UC biopsy specimens^8,10^. Neutrophils and monocytes exhibit similarities in gene expression, so there could be overlap in the annotation of neutrophils and monocytes. Here, the monocyte population exhibited higher *HLA-DRA, HLA-DRB1,* and *VCAN,* while the neutrophils expressed higher *CSF3R, FCGR3B, S100A8, S100A9, and TREM1 (***Supplementary Figure 8C**. To investigate this interaction further, we calculated the neighborhood enrichment z-scores between IAF and monocytes for each individual core and averaged the scores per patient. This revealed low enrichment scores in HC, with multiple instances where z-scores could not be calculated due to low or no abundance (**Figure 5D**). UC PRE biopsies showed a trend (p value 0.07) toward increased z-scores for IAFs and monocytes, consistent with the increased abundance and proximity of IAFs and monocytes in colitis. Spatial scatter plots of these subsets within representative HC and UC PRE samples confirmed these findings (**Figure 5E**). Taken together, we define an IAF-monocyte cellular network that increases in abundance and spatial proximity in patients with active UC.

Finally, we examined the biopsies from UC patient’s pre-treatment with vedolizumab, stratified by subsequent treatment response, to assess the association of spatial cellular network with treatment outcomes. We performed a pseudobulk DEG analysis, stratifying the UC PRE group into responders (UC PRE R) and non-responders (UC PRE NR) to VDZ (**Supplementary Figure 9, Supplementary Table 6**). Consistent with previous findings^9^, our Xenium dataset identified response-associated genes linked to IEC crypts, including *AGR2*, *PIGR*, and *SPINK4*. Additionally, we observed an increase in IAF and MNP-associated genes, such as *MMP1*, *MMP3* for IAF and *FCER2, CD1C* for the MNP, in non-response (**Supplementary Figure 9A-C**). Moreover, the Xenium dataset enabled the detection of differences in genes associated with B cells (*CD19, MS4A1, BANK1* and *SELL*) in UC PRE NR versus UC PRE R (**Supplementary Figure 9A-B**). These B cell genes were located spatially in GALT aggregates (**Supplementary Figure 9C**). This raised the possibility that VDZ non-responders have higher levels of GALT pre-treatment compared to responders, supporting recent work showing that VDZ reduces GALT in responders after treatment^21^. We then performed GSEA of external bulk transcriptomic data using our Xenium-defined gene sets from VDZ responders and non-responders. GSEA validated our Xenium VDZ response signature, comprised largely of epithelial genes (**Supplementary Figure 9D**). Interestingly, the overall Xenium VDZ non-response signature was not validated by GSEA in this external dataset. We then divided the Xenium VDZ non-response signatures into two separate signatures, one comprised of genes from IAF, monocytes, and neutrophils, and the other comprised of genes enriched in GALT aggregates, including B cell, DC, and S4 fibroblast landmark genes. The IAF-monocyte-neutrophil signature was enriched in VDZ non-responders by GSEA in the external dataset, but the GALT-B-DC-S4 fibroblast signature was not, suggesting that there may be some heterogeneity in VDZ non-responders. These data demonstrate the potential for Xenium to identify pre-treatment response and non-response signatures using archived clinical FFPE specimens.

## DISCUSSION

To identify the optimal spatial transcriptomics platform for analyzing archived, clinical FFPE mucosal biopsies of non-IBD patients and patients with UC, we compared three commercially available platforms on the same FFPE TMA. The Vizgen MERSCOPE was not able to analyze these samples due to difficulties with tissue clearing. Comparing 10X Genomics Xenium and Nanostring CosMx, the Xenium detected more transcripts and genes per cell for overlapping genes with significantly lower non-specific probe detection. The smaller custom Xenium panel, optimized for mucosal biopsies of patients with IBD, enabled the identification, annotation, and quantitation of more cell subsets and identified more differentially expressed genes across cell subsets and disease states. The default nuclear expansion cell segmentation with Xenium inflated the cell sizes of a small percentage of cells but identified comparable median cell area to other algorithms with minor impact on the overall results. Using this improved spatial pipeline, we mapped transcriptionally distinct fibroblast subsets to discrete spatial locations, and we identified increased abundance and trends toward increased spatial proximity of IAFs and monocytes in patients with UC. We also identified signatures associated with response or non-response to VDZ, highlighting IAF-monocyte networks and B cell-GALT abundance in non-responders as well as IEC gene sets in responders. Spatial transcriptomics preserves cell subset frequency and transcriptional states with high fidelity, and these methods can facilitate robust tissue profiling using archived FFPE.

Several recent studies have examined imaging-based spatial platforms on tumor tissue, and the results are largely consistent with this study^14–16^. One study found higher sensitivity and broader dynamic range with Xenium for FFPE prostate cancer samples^15^. Another study of FFPE tumor blocks originating from breast, colorectal, lung, ovary, bladder, and lymphoma similarly observed better gene clustering, fewer false positives, and better correlation with whole transcriptome approaches for Xenium as compared to CosMx^16^. A third study of 7 tumor types and 16 normal tissues found higher transcript counts, higher specificity, and lower false discovery rates for Xenium compared to CosMx, and showed that MERSCOPE was the most dependent on RNA integrity^16^. The default Xenium cell segmentation consistently inflated cell sizes across all these studies, but the overall impact on performance was minor, consistent with our findings. It is important to note that the preferred spatial transcriptomics approach may depend on the upstream processing and tissue type. For example, one recent benchmarking study using fresh-frozen mouse brain samples demonstrated superior performance with Vizgen’s MERSCOPE over the Xenium platform^22^. Therefore, considering the tissue type and sample processing protocol will be critical for choosing a spatial transcriptomics platform.

It is also important to acknowledge that some of the improvements in cell annotation we observed with Xenium were due to the customized panel for IBD, which included key landmark genes for our cell subsets of interest. This focused panel allowed us to map transcriptionally distinct fibroblast subsets to discrete spatial locations, for example, confirming S2 as sub-epithelial crypt-associated fibroblasts, S3 as submucosal fibroblasts, and S4 as follicular stromal cells co-localizing with lymphoid aggregates in GALT. Although it would be appealing to employ an unbiased universal panel across tissues, our results suggest that an optimized panel for a specific tissue type and disease state will capture more cells and genes of interest with improved performance. In IBD, there are ample reference tissue scRNA-seq datasets to guide panel design^1–4,6,7,9^, but other tissues and disease processes may benefit from initial studies using larger, unbiased panels or whole transcriptome approaches. While we observed excellent cell subset identification using a smaller, custom IBD panel, this focused panel may not capture the breadth of differentially expressed genes offered by larger panels or whole transcriptome methods. Due to optical crowding with imaging-based spatial transcriptomics platforms, there will likely continue to be a trade-off between the signal for each gene and the size of the panel. Therefore, multiple complementary spatial transcriptomics approaches might be necessary to adequately capture the transcriptional diversity of cell subsets and disease states in FFPE tissue. Whether focused or whole transcriptome methods are used, the transcript counts for these spatial methods are generally orders of magnitude lower than scRNA-seq, which could limit downstream differential gene expression analyses. Therefore, bulk RNA-seq and scRNA-seq will continue to be critical for dissecting tissue inflammation and informing spatial transcriptomics datasets.

The identification of both B cell abundance in GALT and IAF-monocyte networks in UC biopsies, including some VDZ non-responders, complements recent studies regarding UC and the effects of VDZ. Comprehensive flow cytometry analysis of mucosal biopsies identified a significant reduction of CD1c^+^ (BDCA1^+^) type 2 conventional dendritic cells (cDC2s) in UC patients on VDZ^23^. We subsequently reported a shift of α4β7^+^ conventional dendritic cells (cDCs) from the tissue to the circulation in UC patients on VDZ^9^. Interestingly, another group reported that VDZ reduced naïve B and T cells and gut-homing plasmablasts, culminating in GALT attrition in VDZ responders^21^. Given the central role of cDCs in priming naïve T and B cells in lymphoid tissue, VDZ may reduce GALT by markedly inhibiting migratory cDCs from trafficking to the colon. Here, we identified higher B cell abundance and *CD1C* expression in some VDZ non-responders pre-treatment by Xenium. Given that both B cells and CD1c^+^ cDCs localize to GALT, these data imply that higher pre-treatment GALT area may be associated with VDZ non-response, although the GALT signature in VDZ non-responders was not validated in an external transcriptomic dataset. Recruitment of neutrophils and monocytes to inflamed colon tissue by IAFs has also been associated with UC and non-response to VDZ^8,9^. In this study, we also observed an increased abundance and trend toward proximity of IAFs and monocytes in UC compared to HC, confirming prior associations of this cellular network with colitis. Neutrophils were also increased in abundance in patients with UC in this study, and given similarities in gene expression, there is likely some overlap in the annotation of neutrophils and monocytes using spatial transcriptomics. Finally, we identified pre-treatment spatial transcriptomics signatures in the epithelial compartment associated with a response to VDZ, suggesting a potential mechanism for mucosal healing. Namely, VDZ responders may have IECs that are poised to regenerate the epithelium once inflammation is suppressed.

There are important limitations of this study to consider, and several caveats regarding spatial transcriptomics in general. First and foremost, we compared healthy and inflamed colon biopsies and VDZ responders versus non-responders using a small number of patients. Future longitudinal spatial transcriptomics studies will need to include more patients on various therapies to identify signatures predictive of response or non-response. We also focused exclusively on archived human FFPE colon tissue, which is a unique tissue type. These endoscopic mucosal biopsies are placed directly in formalin within seconds of collection, they permeabilize easily due to their small size, and embedding is typically completed within a day or a few days. Thus, our findings may not extrapolate seamlessly to other specimens that vary in size and fixation times. As noted above, several studies have examined imaging-based spatial platforms on tumor and normal tissue, and the results largely align with our study^14–16^, but certain tissues or disease processes may be captured differently by various platforms. Moreover, additional gene sets would be required to adequately profile ileal samples. Another fundamental limitation of spatial transcriptomics is that annotating cells exclusively using transcriptional signatures has inherent drawbacks. For example, confidently differentiating naïve, effector, regulatory, and memory T cell subsets would be markedly improved with simultaneous spatial proteomic surface marker analysis. Similarly, the identification of polymorphonuclear cells (PMNs) and IELs would be greatly enhanced with expert concurrent annotation of H&E images by a pathologist, rather than relying solely on landmark gene expression. Spatial transcriptomics studies are also limited by sampling variability. For example, here we analyzed 4μm-thick sections of cores from 2-3mm biopsies meant to reflect the inflammatory status of an organ approximately one meter long. Adequate tissue sampling will be critical to consider in all future spatial transcriptomics studies.

In summary, we compared three commercially available platforms and optimized a custom spatial transcriptomics panel for analyzing archived FFPE colon mucosal biopsies from patients with or without colitis. Our custom IBD-specific Xenium panel detected more transcripts and genes per cell for overlapping genes, identified more unique cell subsets, quantitated more significant changes in cell frequency, and detected more differentially expressed genes across disease states. This approach yielded improved annotation and spatial analysis of fibroblast subsets and uncovered IAF-monocyte neighborhood enrichment in inflamed colon biopsies. This study adds to the growing body of literature identifying critical interactions among MNP, neutrophil, and fibroblast subsets in inflamed tissues across multiple diseases. We also identified increased B cell frequency in UC biopsies compared to HC, particularly in VDZ non-responders pre-treatment, demonstrating the ability of spatial transcriptomics to profile alterations in GALT across disease states. The capability of generating spatial transcriptomics data from routinely collected, archived clinical specimens with subcellular resolution could enable transformative translational research. Spatial transcriptomics could accelerate the identification and validation of candidate biomarkers and nominate novel therapeutic targets in affected tissue for a variety of inflammatory disorders.

## METHODS

### Study approval

The study was conducted according to the Declaration of Helsinki principles and was approved by the Institutional Review Board of the University of California, San Francisco (19-27302). Written informed consent was received from participants prior to inclusion in the study.

### Study participants and biospecimen collection

For this study, both prospectively collected and retrospectively retrieved archived formalin-fixed paraffin-embedded (FFPE) samples were used. Healthy control (HC) patients refer to those without known or suspected inflammatory bowel disease (IBD) who were undergoing routine colonoscopy or sigmoidoscopy for various indications, such as colorectal cancer screening. HC biopsies were prospectively collected, placed in 10% formalin in 5 mL tubes for approximately 24h, then washed with PBS three times, and then transferred to 30%, 50%, and finally 70% ethanol, and they are stored in 70% ethanol for up to a month prior to paraffin embedding. All UC specimens were retrospectively retrieved. For retrospective retrieval, study subjects were identified by querying the electronic medical records of patients previously seen by UCSF Gastroenterology with existing archived specimens. This was followed by obtaining written informed consent and approval. Baseline demographic and clinical information about the study participants are provided in **Supplementary Table 1.** The CosMx dataset from this cohort was previously published and re-analyzed for this study^9^. Xenium and MERSCOPE analyses were acquired for this study as detailed below. We have consent to publish de-identified patient demographics including age at the time of sample collection, sex, diagnosis, and medical center. Demographic options were defined by the investigators, and participants chose their classifications. Biopsy samples were categorized based on the level of inflammation observed endoscopically: non-inflamed (score=0), mildly inflamed (score=1), moderately inflamed (score=2), or severely inflamed (score=3). Samples were assigned unique identifiers before biobanking.

### Histology FFPE tissue microarray (TMA) construction

Tissue microarrays were constructed from FFPE blocks by Pantomics, with 1.1–1.5 mm cores. The IBD TMA contained multiple cores from 9 prospectively-collected HC and 11 retrospectively-retrieved UC patients. For UC patients, cores were obtained both before and after treatment with Vedolizumab. Core areas were selected based on individual H&E staining of each block, with representative areas chosen per core. Core samples from the same patients had at least duplicate cores from different locations, although some cores were not within the fiducials and final scanned area. The recipient TMA block was sectioned with a clearance angle of 10° and a thickness of 4 μm along the width of the block and was used for H&E staining. The TMA block was stored at 4 °C. Freshly cut TMA sections were prepared before each experiment, according to the recommended protocol for each spatial transcriptomic assay performed.

### CosMx spatial transcriptomics tissue processing, staining, and imaging

FFPE TMA processing, staining, imaging, and cell segmentation were performed as previously described and published^24,9^. The time from sectioning to CosMx SMI analysis was 9 days. For this analysis, the raw data files were re-loaded into our preprocessing and technical performance comparison Jupyter notebooks to ensure consistent analysis across spatial transcriptomics platforms.

### Xenium spatial transcriptomics tissue processing, staining, and imaging

Xenium sections were prepared following the Xenium “In Situ-Tissue Preparation Guide protocol” (10X Genomics, Demonstrated Protocol, CG000578-Rev C). Briefly, 4 µm sections of the FFPE TMA block were sectioned, and shortly kept afloat in an RNAse-free water bath at 42°C. They were then carefully transferred from the water bath to the marked sample area of the Xenium slide (PN-1000460). To remove excess of water, the slides were placed upright in a drying rack at room temperature for 30 minutes and then baked for 3 hours at 42°C. Thereafter, the slides were stored in a vacuum-sealed bag at room temperature until they were ready for processing. The time from sectioning to Xenium analysis was 2 days for replicate 1 and 9 days for replicate 2. The Xenium slides were processed at UCSF according to the Xenium “In Situ for FFPE-Deparaffinization and Decrosslinking protocol” (10X Genomics, CG000580-Rev C). The slides were incubated at 60°C for 2 hours and then cooled down at room temperature for 7 minutes. After cooling, the slides were immersed first in xylene, then ethanol, and lastly, nuclease-free water to deparaffinize and rehydrate the tissue. Post rehydration, slides were assembled into the Xenium cassette (10X Genomics, PN-1000566) and the tissue sections were decrosslinked using the Xenium Slides and Sample Prep Reagents Kit (10X Genomics, PN-1000460). Decrosslinking involved a thermocycler incubation at 80°C for 30 minutes and then 22°C for 10 minutes with the decrosslinking buffer. The slides were washed multiple times with PBS-T and, subsequently, were hybridized following the Xenium “In Situ Gene Expression User Guide” (10X Genomics, CG000582 Rev D). Tissue slides were incubated at 50°C overnight with a customized gene panel, which included 290 genes (**Supplementary Table 2**). The next day, unbound probes were removed by washes with PBS-T and a thermocycler incubation at 37°C for 30 minutes with the post-hybridization wash buffer. After the removal of unbound probes, the junction between the RNA with its hybridized probes was ligated using the ligation mix in the thermocycler at 37°C for 2 hours. The probe-RNA product was then amplified at 30°C for 2 hours with the Amplification Master Mix to generate multiple gene-specific barcodes. Post amplification, the slides were washed and quenched with autofluorescence quencher to reduce signal noise. Lastly, before loading the Xenium cassette in the machine, nuclei were stained with DAPI. Xenium slides were imaged using the Xenium Analyzer following the guidelines provided in the Xenium Analyzer User Guide (10X Genomics, CG000584-RevB). The decoding modules A and B were transferred from 20°C to 4°C a night prior to the instrument run. On the day of the instrument run, first, the buffers (Wash Buffer A, Wash Buffer B, Xenium Instrument Wash Buffer, and Xenium Probe Removal Buffer) were prepared and loaded into the machine. Then, the decoding reagent plates, pipette tip rack, extraction tip, and the wetting consumable were loaded. Lastly, the Xenium cassettes containing the slides were loaded into the Xenium Controller machine. After successful loading, the machine took about an hour for a sample scan. From the output of the scan on the machine screen, we selected FOVs containing the tissue cores and started the run. Nucleus boundaries were identified using a nucleus segmentation algorithm applied to the nuclei-stained (DAPI) morphology image, and cell boundaries were determined by expanding the nucleus boundaries by 5 µm or until they intersected with another cell. The cell-specific and transcript-specific metadata, found in the cell summary and transcript output files, respectively, included several cell segmentation-related metrics: cell area (µm^2^) and nucleus area (µm^2^) for cells, and overlap with the nucleus (yes/no options) and nucleus distance (µm) for transcripts. After the Xenium run was completed, all data were stored in an organized output directory on the Xenium instrument. Data were downloaded and we used the cell summary file (cells.csv.gz) and the cell-feature matrix file (cell_feature_matrix.h5), which included the x and y coordinates for each individual cell. The transcript file (transcripts.csv) was used for the QC transcript-level comparisons.

### MERSCOPE spatial transcriptomics tissue processing, and staining

MERSCOPE slides were prepared according to “MERSCOPE User Guide-Formalin-Fixed Paraffin-Embedded Tissue Sample Preparation” (91600112 Rev C). Briefly, 4 µm sections of the FFPE TMA block were sectioned, and place onto the center of the MERSCOPE FFPE slide (PN 20400100). Then, slides were transferred to a drying rack to remove excess water. Slides were dried at 55°C for 15 minutes and then at room temperature until no visible water droplets were present. Dry slides were stored in a 60-mm petri dish and sealed with parafilm at −20°C until further processing. The slides were processed within 6 weeks of sectioning. Before running the samples with the full gene panel, one of the MERSCOPE slides was used to run sample verification kit (91600004 Rev D), targeting a housekeeping gene EEF2 in Homo sapiens to verify RNA quality, assess background noise, and optimize protocol conditions. The assay using the full customized gene panel was then conducted. The MERSCOPE slide was equilibrated to room temperature for 30 minutes and underwent deparaffinization and decrosslinking steps. Subsequently, a pre-anchoring treatment (PN 20300116; PN 202300113) was performed at 37°C for 2 hours, followed by a cell boundary staining (PN20300100) with a primary staining solution at room temperature for 1 hour. Then, the MERSCOPE slide was washed three times with 1X PBS and incubated with a secondary staining solution at room temperature for 1 hour. After this step, samples were incubated with formamide wash buffer (PN 20300002) at 37°C for 30 minutes and subsequently incubated with RNA anchoring buffer (PN 20300117) at 37°C overnight. Following overnight incubation, another formamide wash was conducted at 47°C for 15 minutes and the slide was washed before to proceeding to the gel embedding step (PN 30200004). For resistant FFPE tissues, a digestion step (PN 20300005) was performed at 37°C for 5 hours before incubation in clearing solution (PN 20300114) for 13 days at 37°C to ensure tissue transparency, instead of the recommended 7 days in clearing solution, based on guidance from the manufacturer. Tissue transparency was never fully achieved. Consequently, MERSCOPE slides underwent autofluorescence quenching using MERSCOPE photobleacher (PN 10100003). Next, clearing solution was removed and slides were washed and incubated with formamide wash buffer at 37°C for 30 minutes before incubating with the customized 280-plex gene panel (**Supplementary Table 2**) in a humidified 37°C cell culture incubator for at least 36 hours. Finally, a post-encoding probe hybridization wash was completed through two incubations with formamide wash buffer at 47°C for 30 minutes each. Then, the slide was washed twice with sample wash buffer and incubated with DAPI and PolyT staining reagents at room temperature for 15 minutes. The slide was then assembled into MERSCOPE flow chamber and loaded on the MERSCOPE instrument for imaging following the User guide (91600001 Rev G) for proper configuration and data acquisition. Segmentation parameters were selected and image processing was run using the Cellpose segmentation algorithm to produce single-cell outputs^25^. The cell-feature matrix file (cell_by_gene.csv), cell metadata file (cell_metadata.csv), and cell and transcript location information file (micron_to_mosaic_pixel_transform.csv) were used for our data processing analysis.

### Technical performance comparisons among spatial transcriptomics platforms

Our technical QC comparison between Xenium and CosMx platforms utilized transcript data, filtered to only include high-quality transcripts that passed platform-specific filters (Xenium platform phred-scaled quality value (Q-Score) > 20; All CosMx QC flags = ‘Pass’) and our cell-level filtering schema, which included filtering out cells with <50 counts and <10 genes and filtering out genes with <1 count and detected in <10 cells.

The sources of the data values used for various QC comparisons are detailed below. To quantify cellular location classification for transcripts, we used the Xenium transcript metadata ‘Overlaps Nucleus’ metric (yes/no options) and the CosMx transcript metadata ‘Cellular Compartment’ metric (0, cytoplasm, membrane, and nuclear options). Post-filtering, we had 0 transcripts with 0 values. For the cellular location classification plot, we grouped cells as nuclear and non-nuclear for both platforms. Cell area was compared across platforms using µm^2^ values. For Xenium, the cell area values were already provided in µm^2^. However, for CosMx, the cell area values were initially in pixels and were converted to µm^2^ by multiplying each pixel value by 0.0144, as 1 pixel equals 0.12 µm. Negative probes per cell were quantified using the ‘Control Probe Counts’ cell metadata column in Xenium and ‘nCount NegProbes’ cell metadata column in CosMx. Signal over background metrics were calculated by dividing the mean expression of each gene within cells by the mean negative probe counts per cell. Block age was measured from the time of tissue collection until the assay was conducted. As a specificity measure, we calculated mutually exclusive co-expression rate (MECR) values for each dataset^22^. We used lineage-specific genes that are highly expressed in our cell types of interest within the Xenium, CosMx, and our previous scRNA-seq dataset performed on colon biopsies collected from HC and UC patients (GSE250487). Ideally, these lineage-specific genes should only be expressed within their corresponding cell types. The MECR metric quantifies the detected co-expression rate of two lineage-specific genes in individual cells, normalized for the abundance of each gene. MECR values were calculated for each gene pair by dividing the fraction of cells that express both markers by the fraction of cells that express at least one of the markers.

### Data preprocessing and annotation

We developed and optimized a standard preprocessing pipeline using the Python packages scanpy, squidpy, and anndata, tailored for use with CosMx, Xenium, and MERSCOPE data^26,27^. We created spatial data objects in Python for each data type using cell-level gene expression, metadata, and spatial location information. Low quality transcripts (as qualified by a Q-score < 20) were automatically removed from the Xenium cell-level data before processing steps. For the CosMx and Xenium data, we filtered out cells with <50 counts and <10 genes and genes with <1 count and detected in <10 cells. As the MERSCOPE dataset had very few cells passing QC, we tweaked our filtering schema to remove cells with <10 counts instead of <50 counts. The other filtering criteria remained the same. After filtering, we proceeded to normalize and log-transform the data and regress out unwanted sources of variation, specifically the number of transcripts per cell and the number of unique (gene) transcripts per cell. Next, we scaled the data and ran a Principal Component Analysis (PCA). We computed a neighborhood graph using n_neighbors=10 and n_pcs=30 (CosMx, Xenium) or n_pcs=14 (MERSCOPE). We embedded the neighborhood graph using UMAP, specifying min_dist=0.2 and spread=1.5 (CosMx, MERSCOPE) or spread=1.75 (Xenium), and clustered using Leiden community detection. We annotated cell clusters by known markers and spatial location, using the Chan-Zuckerberg CELLxGENE tool for refinement^28,29^. The CosMx and Xenium datasets were processed with the same pipeline and settings, as noted above. For the Xenium dataset, some leiden clusters were further divided based on readily apparent, distinct landmark gene expression within subclusters. Cluster 07 was subdivided into neutrophils and monocytes, and the remaining cells expressed high levels of *BANK1, IGHD, MS4A1,* and *CD19* and were combined with the B cell clusters 11 and 16. Cluster 02 and 22 were combined into a coarse endothelial cluster, and then divided into arterial, venous, lymphatic, pericytes and Not Otherwise Specified (NOS) subsets. Cluster 15 was subdivided into S3 and S4 fibroblasts. A LYVE1^+^ macrophage subcluster was annotated macrophage cluster 10. Cluster 24 was subdivided into IAF and plasma cells, and those plasma cells were combined with plasma cell clusters 0, 24, and 30.

### Integrating Xenium replicates

We created spatial data objects in Python for each Xenium replicate data using cell-level gene expression, metadata, and spatial location information. We added prefixes to the cell ID values to distinguish between replicates 1 and 2 and added a ‘batch’ metadata column. We proceeded to filter the datasets separately based on our standard filtering schema (filtered out cells with <50 counts and <10 genes and genes with <1 count and detected in <10 cells). Next, we concatenated the two replicate anndata objects along the observations axis (x-axis; rows) using the anndata.concat function. This process preserves all sub-elements of each object while stacking the observations in an ordered manner. By concatenating along the observations axis, we effectively combined data from different cells (observations) into a single dataset. We then added 15,000 to the x-coordinate value of each cell in replicate 2 to offset its value from those in replicate 1. This adjustment allowed us to visualize the spatial coordinates of the concatenated dataset with replicates 1 and 2 displayed side by side rather than overlapping. After concatenating, we proceeded with the preprocessing pipeline as usual: normalization and log-transformation of the data, regressing out unwanted sources of variation, and scaling the data. We ran a Principal Component Analysis (PCA), integrated the data with Harmony^30^ to account for potential batch effects, and constructed a neighborhood graph using n_neighbors=10 and n_pcs=30. We embedded the neighborhood graph using UMAP, specifying min_dist=0.2 and spread=1.5, and clustered using Leiden community detection. We annotated cell clusters by known markers and spatial location, using the Chan-Zuckerberg CELLxGENE tool for refinement.

### Neighborhood enrichment

We performed neighborhood enrichment analyses for fine annotation cell clusters within the Xenium HC and UC PRE data subsets using functions from the squidpy package. To do this, we constructed a spatial nearest-neighbor graph using Delaunay triangulation, which links cells based on their spatial proximity to each other within a connectivity matrix. We then calculated an enrichment z-score for each pair of fine annotation cell clusters based on cell proximity within the connectivity graph. These analyses were conducted for the Xenium HC and UC PRE data subsets in two ways: (1) by calculating neighborhood enrichment z-scores for each pair of fine annotation cell clusters across the entire dataset (all cores, all patients), (**Supplemental Figure 7A-B**), and (2) by calculating neighborhood enrichment z-scores specifically for our cell types of interest, Fibroblast_IAF and MNP_monocyte cell clusters, within each individual core and then grouping these z-score values for each unique patient (**Figure 5D**).

### Pseudobulk DE genes analysis and Gene Set Enrichment Analysis (GSEA)

Spatial transcriptomic data were used to perform pseudobulk differential expression (DE) analysis using DESeq2^31^. Three distinct Pseudobulk DE gene analyses were conducted: CosMx data comparing UC PRE to HC, Xenium data comparing UC PRE to HC and, Xenium data comparing UC PRE Non-Responders to UC PRE Responders. Samples were stratified by patients. The bulk transcriptomic dataset (GSE73661)^18^, consisting of colonic biopsies from HC and UC patients before and after various biologic treatments, was obtained from the GEO database and utilized for GSEA^32^. For this analysis, the bulk transcriptomic samples were categorized into HC (n=12) and UC PRE (n=43), or UC PRE VDZ R (n=11) and UC PRE VDZ NR (n=9). Data were normalized and z-scored before being processing in the GSEA program (version 4.3.3). HC and UC PRE signatures were defined based on the Pseudobulk DE analysis from Xenium spatial transcriptomic data. For CosMx, only the UC PRE gene signature was identified. Gene signatures obtained from Xenium for UC PRE R, UC PRE NR, IAF-monocyte-neutrophil and GALT-B-DC-S4 fibroblast were explored in the external cohort of patients pre-VDZ stratified by UC PRE R and UC PRE NR. GSEA was then conducted for each gene signature (**Supplementary Table 7**). The number of permutations was set to 1000, with no dataset collapse, using the Affymetrix Human Gene 1.0 ST Array and t-test. For each analysis, a Normalized Enrichment Score (NES) was calculated, and only NES values with a p-value < 0.05 and an adjusted q-value (FDR) < 0.1 were considered significant.

### Statistics

For the comparative analysis among different spatial transcriptomic platforms, the non-parametric method using Mann-Whitney test, two-tailed was utilized. However, for the number of transcripts and unique features per cell per core or FOV, the Kruskal-Wallis test was used to assess differences in population medians across 2-, 3-, 4-, and 5-year old samples. Cell frequencies for Xenium comparing two groups were also assessed using the Mann-Whitney test, followed by a two-stage linear step-up procedure of Benjamini, Krieger and Yekutieli, which adjusts for multiple comparisons by controlling the false discovery rate (FDR). The q-value, representing the FDR-adjusted p-value, was set for discovery at q < 0.1. For DeSeq2 analysis, significance thresholds were defined as log2fc >0.4 or <-0.4 and q < 0.1 and counts threshold of baseMean >500 for Xenium and >400 for CosMx (**Supplementary Table 4-6**). Additional statistical analyses were performed using GraphPad PRISM 10. The ComplexHeatmap R package was used to generate expression z-score heatmaps for DE genes.

## Acknowledgements

MGK is supported by NIH K08 DK123202. MGK holds a Career Award for Medical Scientists from the Burroughs Wellcome Fund. GKF is supported by NIH U01DE028891-01A1, R01AI093615-11, R01DK103735, P30AR070155-05, U01AI168390, R01AI170239, P30 AI027763-31, R01DE032033, and support from the Chan Zuckerberg Initiative, the Bill and Melinda Gates Foundation, Eli Lily, and the UCSF Bakar ImmunoX Initiative. AJC is supported by NIH U01DE028891-01A1, R35CA242447-03, R01HL170038-01 and support from the Melanoma Research Alliance, the California Institute for Regenerative Medicine, Genentech, Eli Lily, and the UCSF Bakar ImmunoX Initiative. This work was also supported by funding from UCSF ImmunoX and the Kenneth Rainin Foundation. Schematics created with BioRender.com. We also acknowledge the Research Core of the UCSF Division of Hospital Medicine. We thank Dr Peng He for his critical feedback on the manuscript. We would also like to acknowledge the valuable contributions from members of the UCSF CoLabs Spatial Transcriptomics Working Group. We thank the study participants for contributing to this research.

## Author contribution statement

Conceptualization: GKF, MGK, AJC; patient recruitment and sample collection: GL, JH, UM, DYO, MGK; sample processing and data acquisition: EM, VJ, CE, GL, JT, JH, WE; cell subset annotation: EM, GL, DEL, JYH; spatial transcriptomics analysis: EM, ML; supervision: GKF, MGK, AJC; funding acquisition: GKF, MGK, AJC; all authors contributed to manuscript preparation.

---

**Supplemental Fig. 1.**
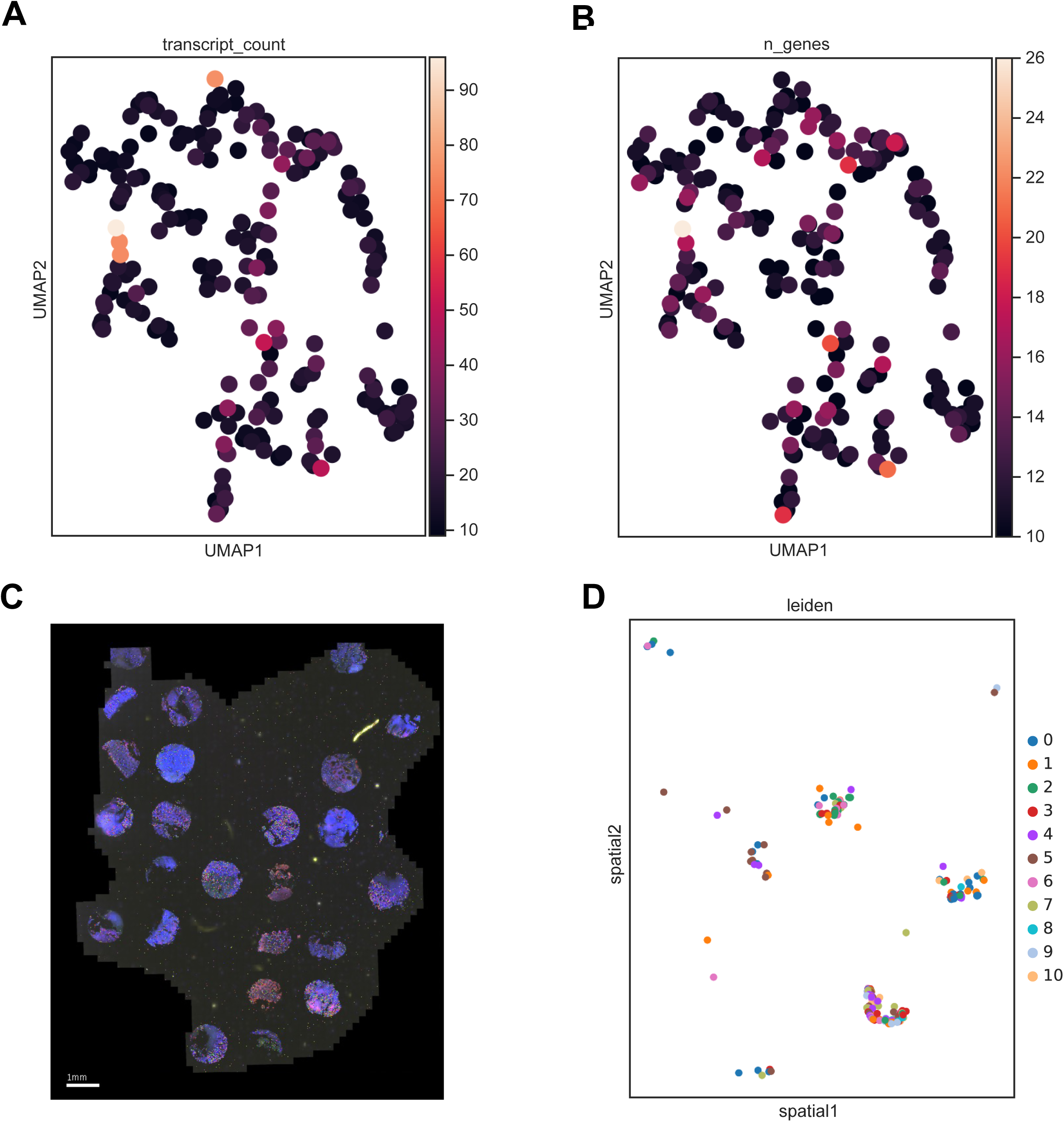
MERSCOPE dataset analysis. (**A**,**B**) UMAP visualization of MERSCOPE dataset (212 cells), highlighting (**A**) transcript count per cell, (**B**) number of genes per cell, and (**C**) area of TMA scanned showing DAPI and detected transcripts (**D**) spatial scatter plot depicting the spatial location of cells in the MERSCOPE dataset in relation to the TMA slide, colored based on leiden clustering.

**Supplemental Fig. 2.**
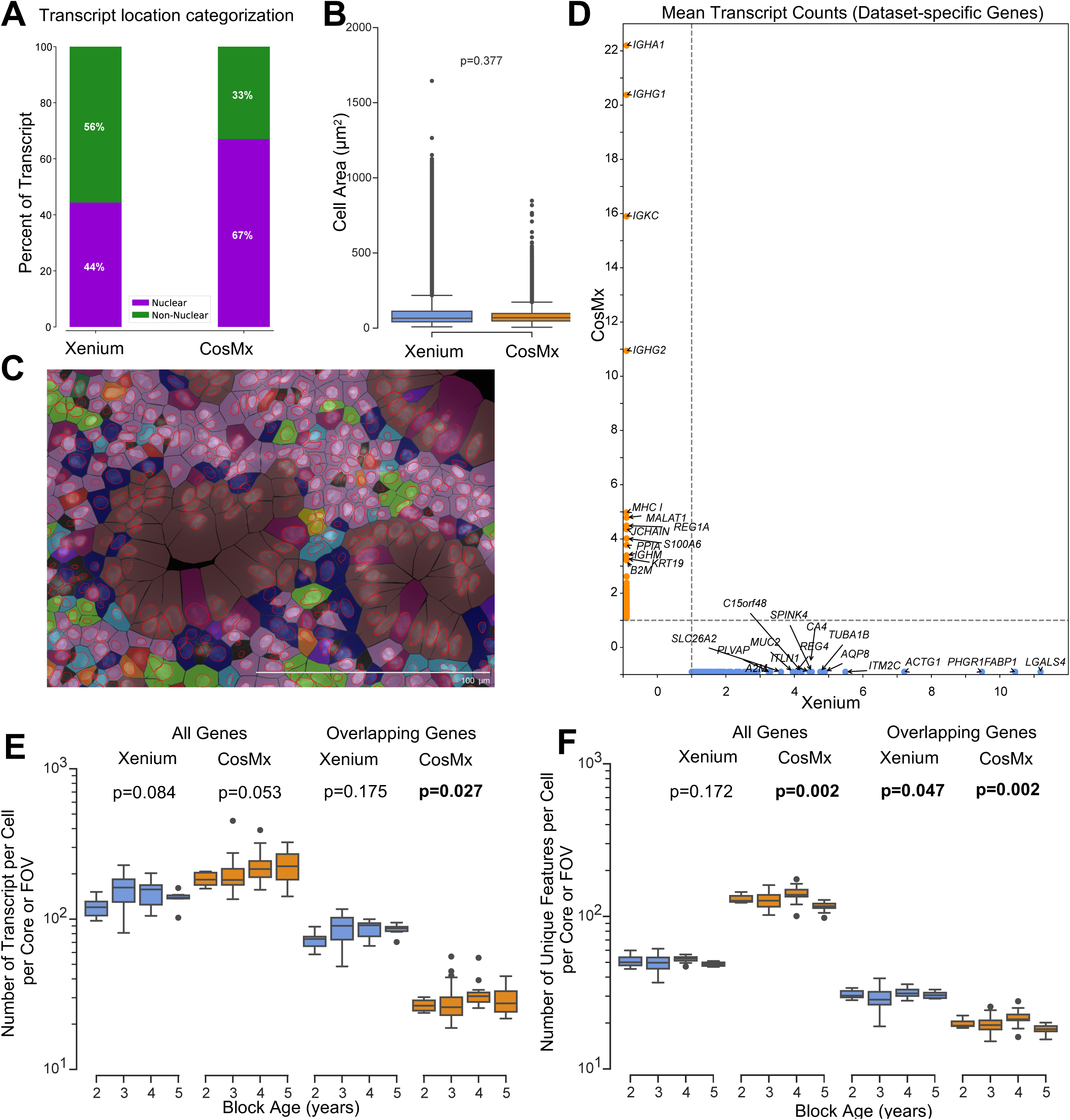
Xenium and CosMx dataset quality control and cell segmentation-related metrics. (**A**) Bar plot comparing the percentage of nuclear versus non-nuclear transcripts within the Xenium and CosMx datasets. (**B**) Cell area plotted in µm^2^ for each cell within the Xenium and CosMx dataset. (**C**) Representative Xenium cell segmentation visualization, displaying DAPI nuclei (white), nuclei outlines (red), cell borders (black), cells are pseudocolored by Xenium fine annotation labels. (**D**) Scatter plot of average transcript count per cell for genes specific to either the Xenium or CosMx (blue for Xenium, and orange for CosMx). (**E,F**) Number of (**E**) transcripts and (**F**) unique features detected per cell per Core or FOV within the Xenium and CosMx datasets, calculated using the complete gene panel for each platform (Xenium, 290 genes; CosMx, 1,000 genes; left) and limited to the 159 overlapping genes across both panels (right). Plotted points are categorized based on the age of FFPE blocks from which they originated. For panels **B, E,** and **F**, box and whisker plots, the band indicates the median, the box indicates the first and third quartiles, and the whiskers indicate minimum and maximum value within the upper/lower fence (upper fence=Q3+1.5xIQR and lower fence= Q1-1.5xIQR), only outlier points are shown; Mann-Whitney, two-tailed tests. Panels E and F compared the difference in population medians between 2-, 3-, 4-, and 5-year-old samples, averaged by core or FOV, with Kruskal-Wallis rank sum statistic (h-statistic) indicated.

**Supplemental Fig. 3.**
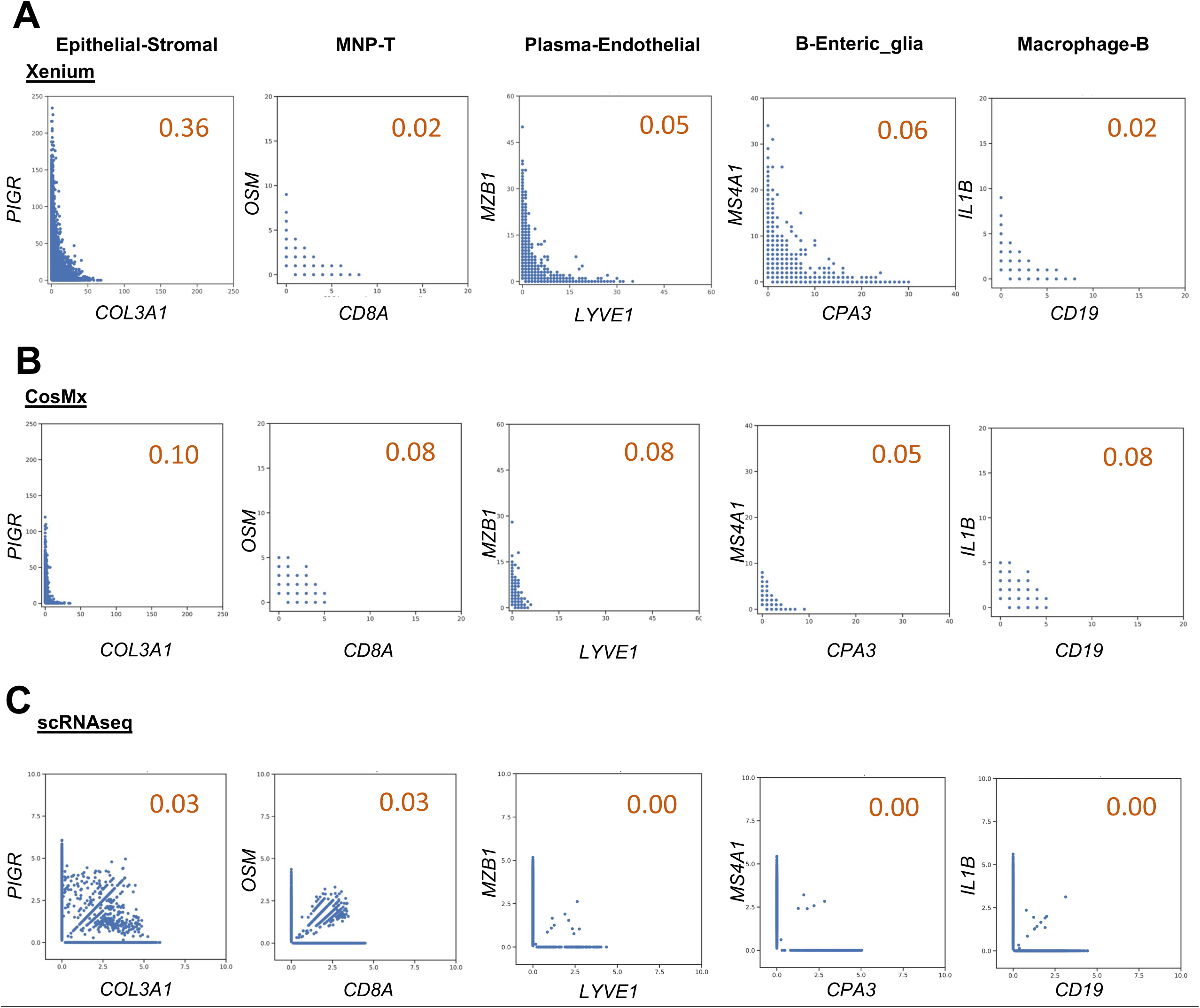
Using mutually exclusive genes to evaluate platform specificity. **(A-C)** Scatter plots of gene expression level (transcript counts per cell) for two representative mutually exclusive lineage-specific genes within (**A**) Xenium, (**B**) CosMx, and (**C**) scRNA-seq datasets generated from colon biopsies collected from HC and UC patients (GSE250487). Genes were selected based on cell types of interest and the overlapping genes between Xenium and CosMx panels. The mutually exclusive co-expression rate (MECR) value is listed for each scatter plot, with lower rates corresponding to greater technology specificity.

**Supplemental Fig. 4.**
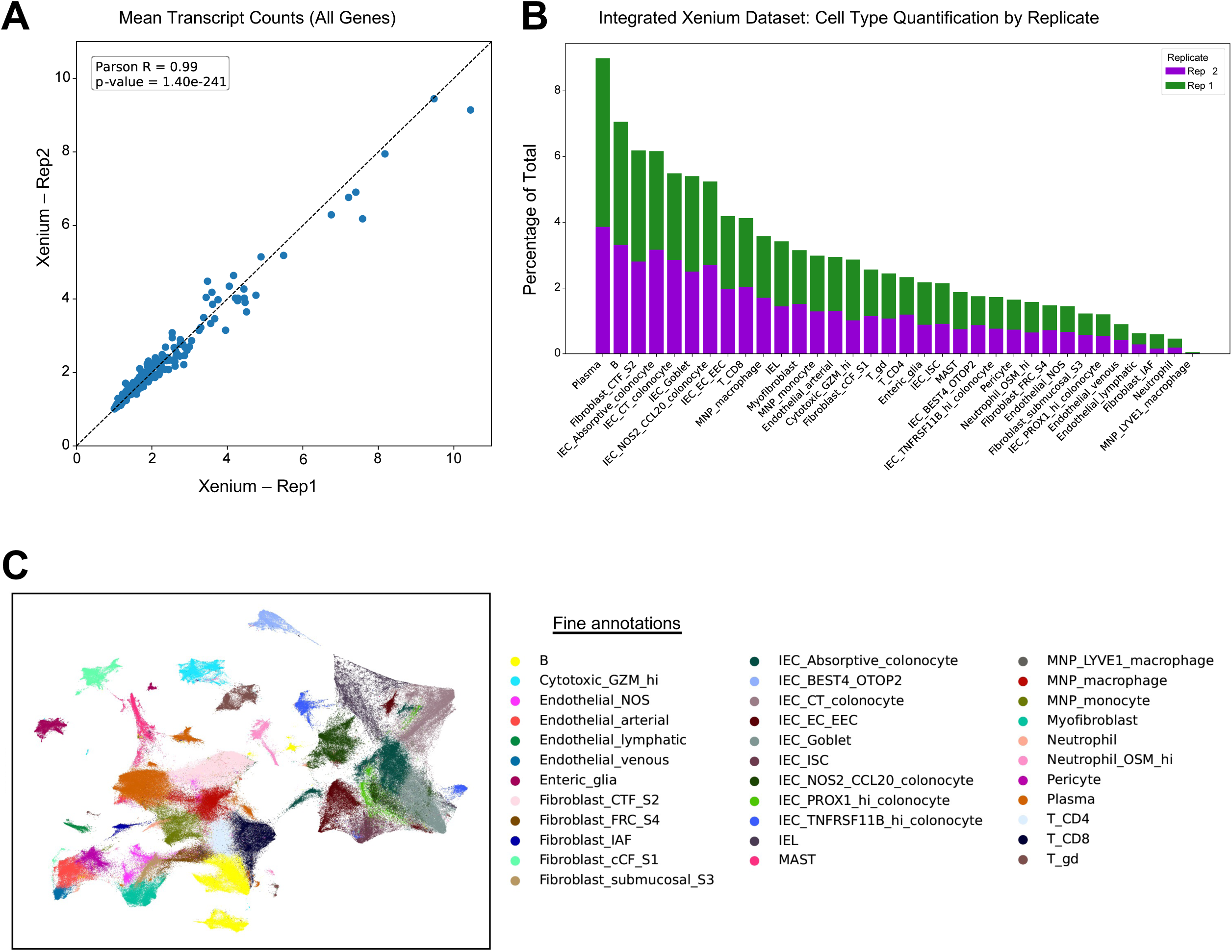
Reproducibility of Xenium replicates. (**A**) Scatter plot showing the gene expression level (transcript counts per cell) for all genes between Xenium replicates. (**B**) Cell frequency as a percent of dataset total for fine annotations within Xenium replicates. (**C**) UMAP visualization of two integrated Xenium replicate runs, colored by fine annotations (582,188 total cells: 313,940 from replicate 1 and 268,248 from replicate 2). Xenium replicate data were obtained from different runs of different sections from same TMA block.

**Supplemental Fig. 5.**
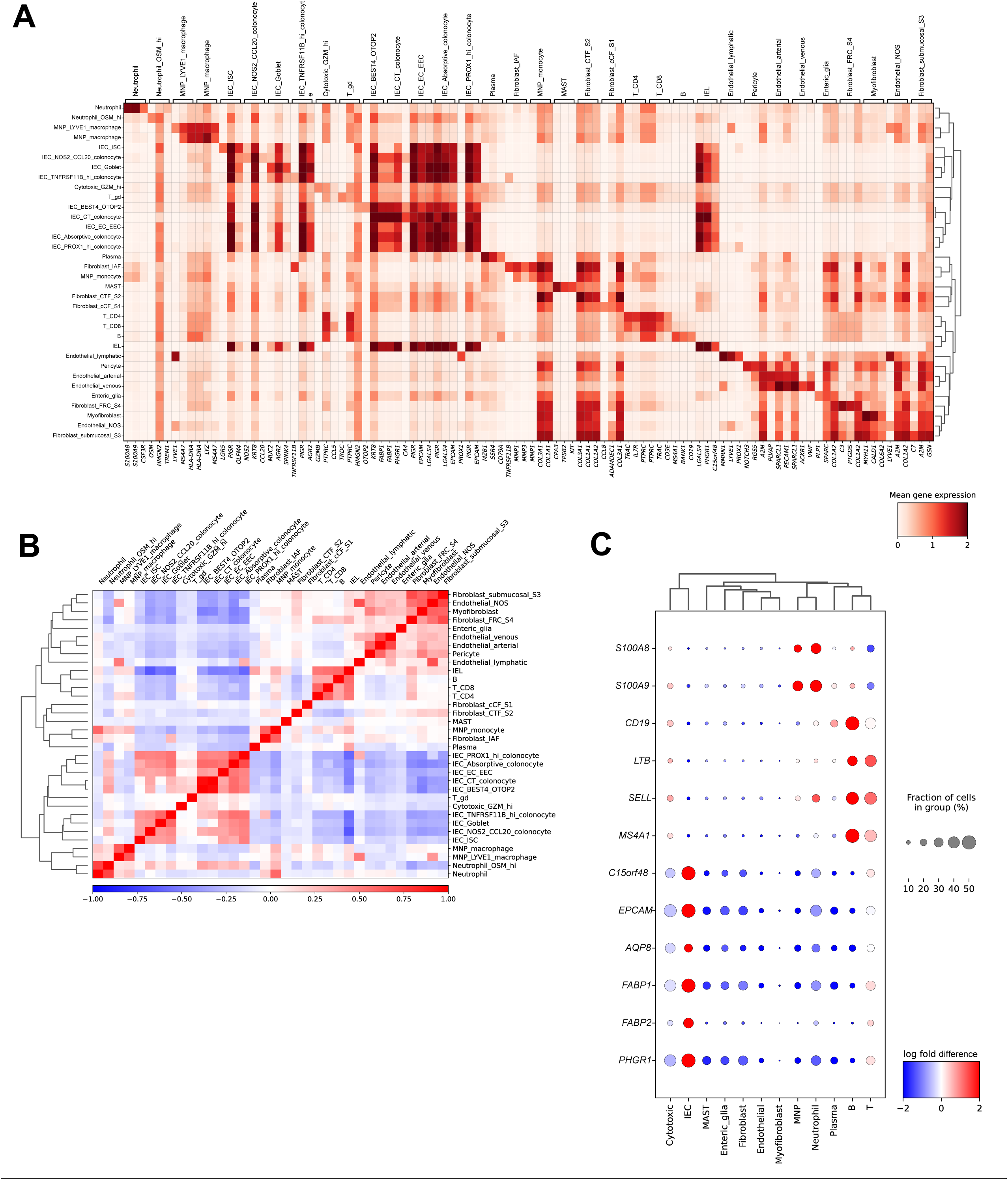
Xenium dataset landmark gene heatmap and correlation matrix for fine annotations, and DEGs between UC and HC. (**A,B**) Fine cell annotation (**A**) heatmap displaying gene expression of the top 3 landmark genes and (**B**) correlation matrix in Xenium replicate 1. (**C**) Dot plot representation of a subset of genes from pseudobulk DEG analysis by coarse annotations.

**Supplemental Fig. 6.**
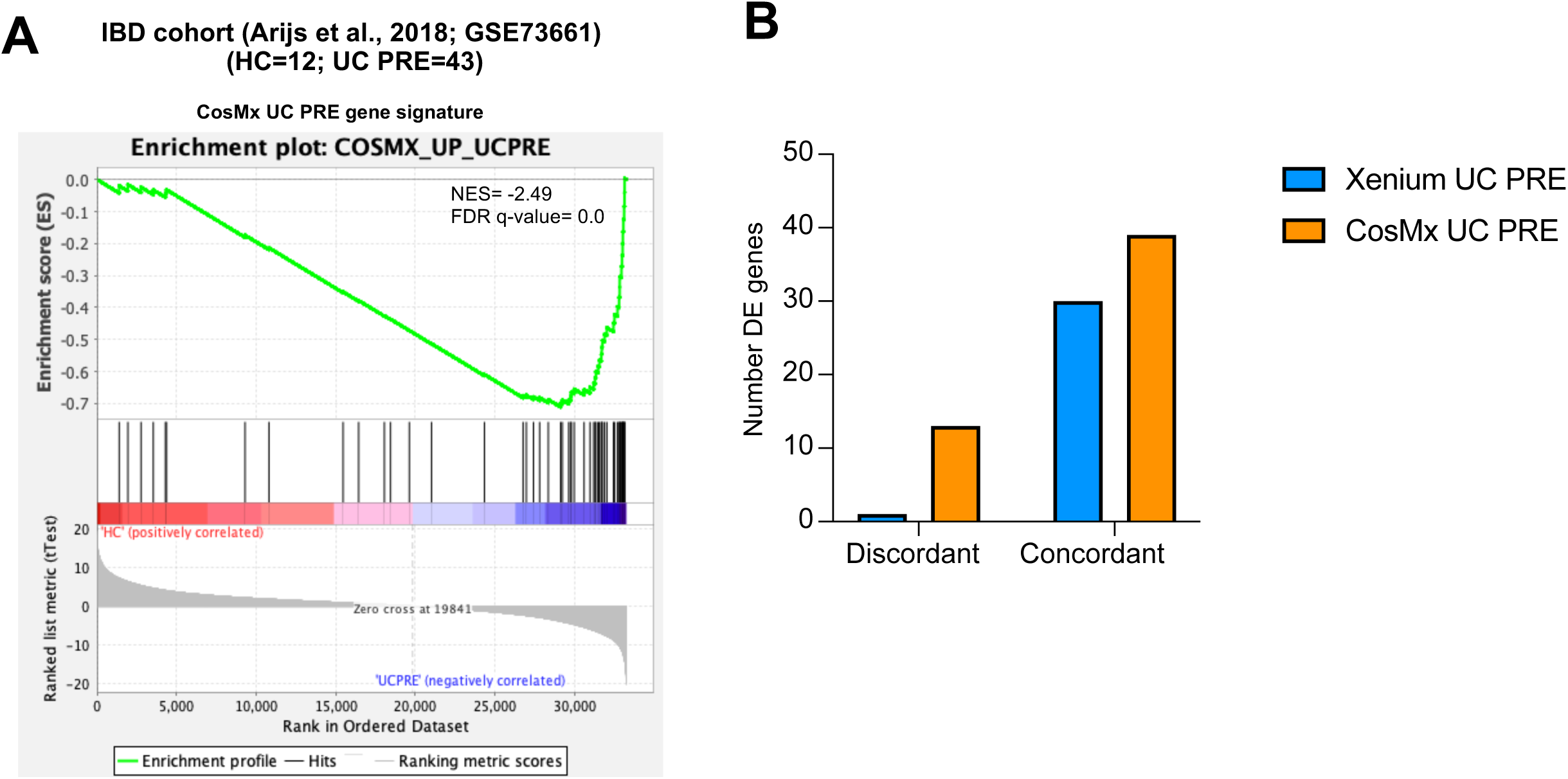
GSEA of an external, publicly-available, bulk transcriptomic dataset using a CosMx transcriptomic signature associated with UC PRE biopsies. (**A**) GSEA of CosMx UC PRE spatial gene signature in an external cohort of patients and relative NES. (**B**) Number of DE genes in the UC PRE signature for Xenium and CosMx that are concordantly or discordantly expressed to the UC PRE patients from the publicly available dataset.

**Supplemental Fig. 7.**
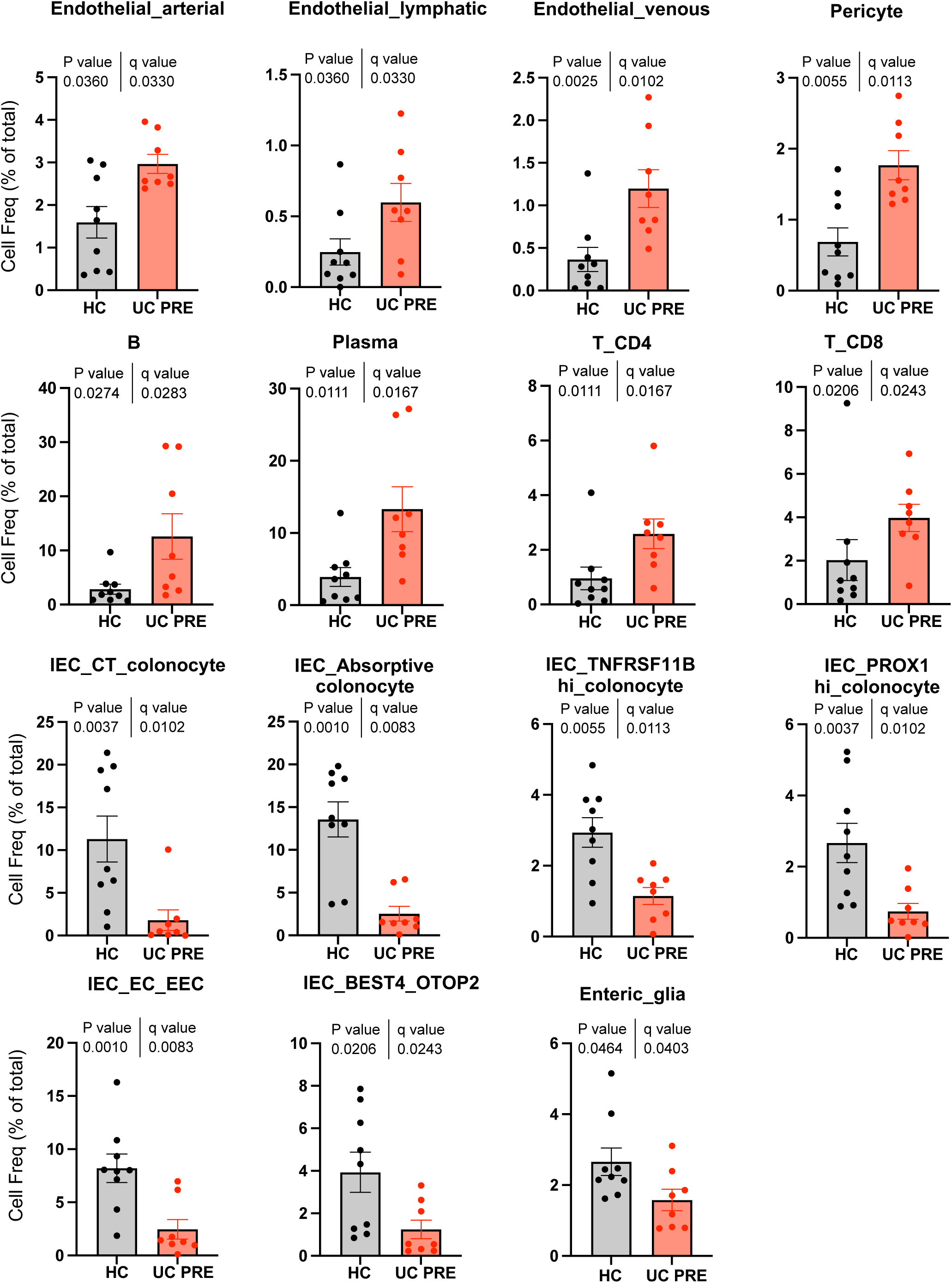
Quantitation of differential cell abundance in the Xenium integrated dataset. Cell frequencies of the indicated subsets comparing HC and UC PRE, each dot represents one patient; Mann– Whitney, two-tailed test with FDR correction; q<0.1 threshold for discovery. Only statistically significant cell subsets are shown with exact p-value and q-value displayed.

**Supplemental Fig. 8.**
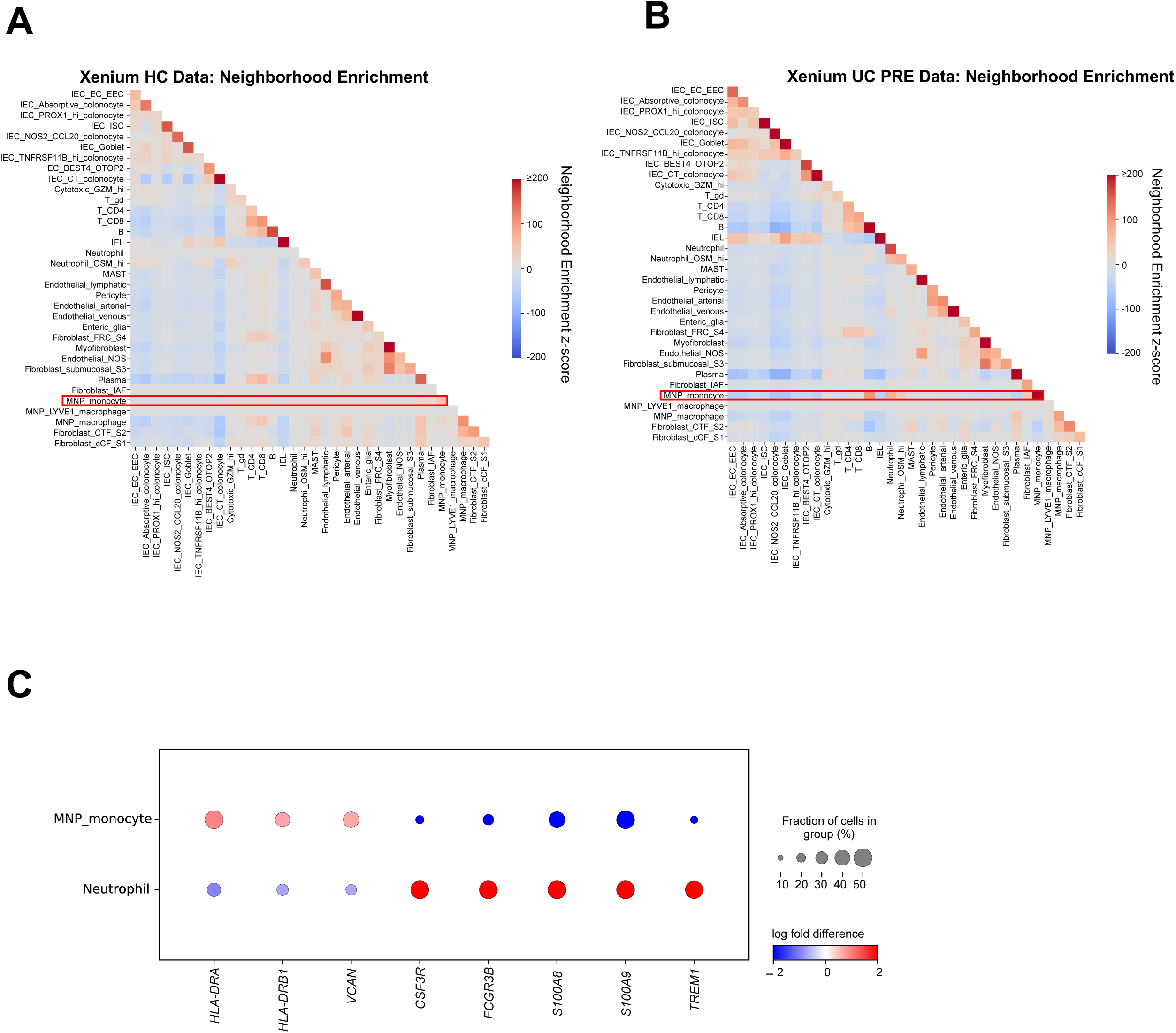
Unsupervised Neighborhood enrichment analysis in Xenium FFPE colon biopsies and dot plot of neutrophil versus monocyte landmark genes. (**A,B**) Heatmaps displaying neighborhood enrichment z-scores for fine annotation cell pairs within (**A**) HC and (**B**) UC PRE biopsies. (**C**) Dot plot representation of landmark genes for the indicated subsets. For panels **A** and **B**, several values exceed the scale for visualization purposes (values greater than 200)..

**Supplemental Fig. 9.**
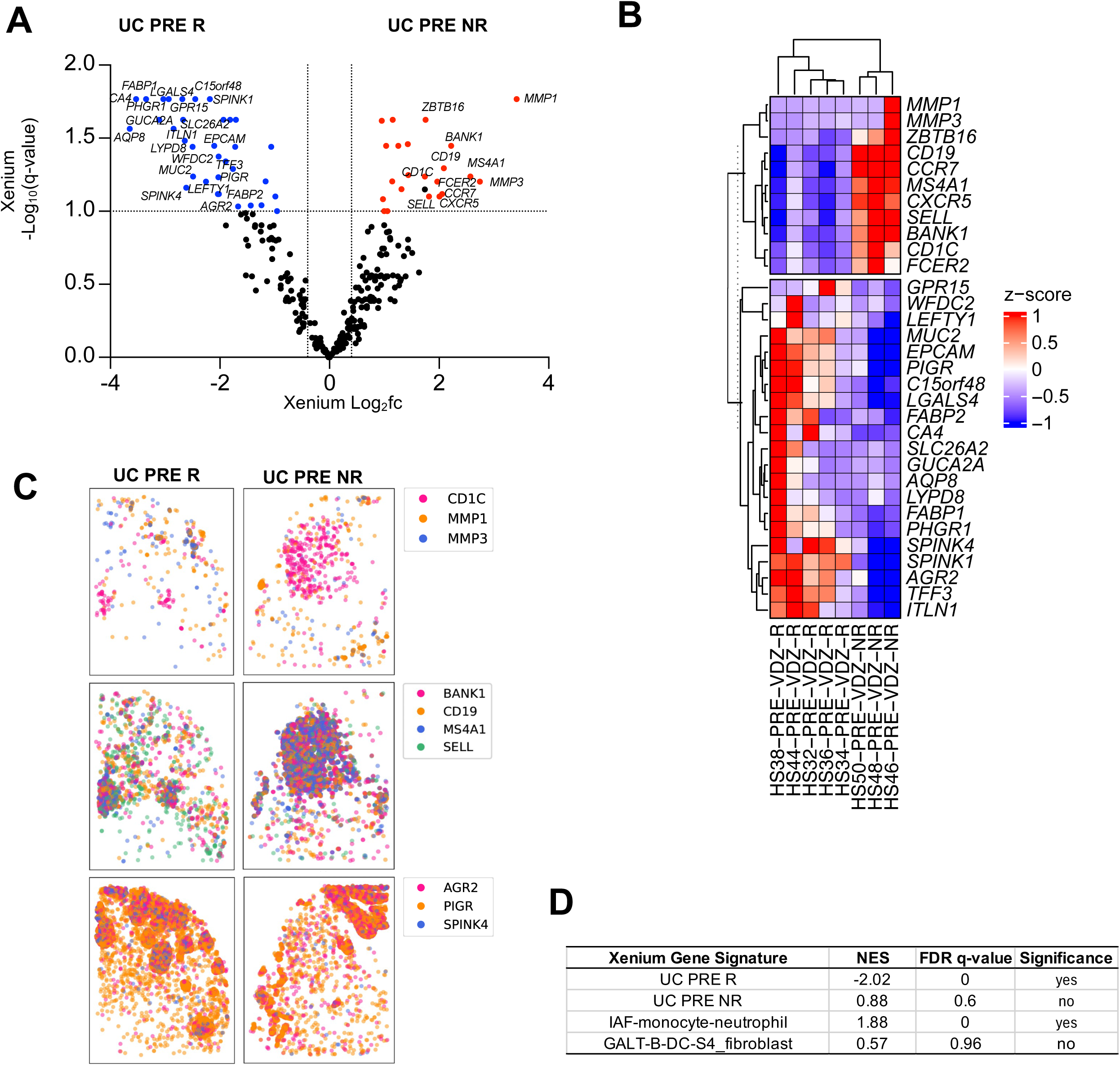
Pseudobulk DE gene analysis comparing UC PRE Non-Responders (UC PRE NR) versus UC PRE Responders (UC PRE R) in the Xenium dataset. (**A**) Volcano plot of pseudobulk DE genes identified by DESeq2 with log2fc >0.4 or <-0.4 and q < 0.1 in UC PRE NR vs UC PRE R. (**B**) Heatmap of expression z-scores for the indicated genes in UC PRE NR (Up/Down) relative to UC PRE R. (**C**) Representative spatial transcript scatter plots highlighting a subset of genes relatively increased in UC PRE R (left) and UC PRE NR (right). (**D**) GSEA of Xenium signatures in external cohort of patients pre-VDZ stratified by UC PRE NR and UC PRE R with relative Normalized Enrichment Scores (NES) and FDR q-value. For panel **B**, some genes are off-scale for visualization purposes, z-score set from −1 to 1.

**Supplementary Table 1.**
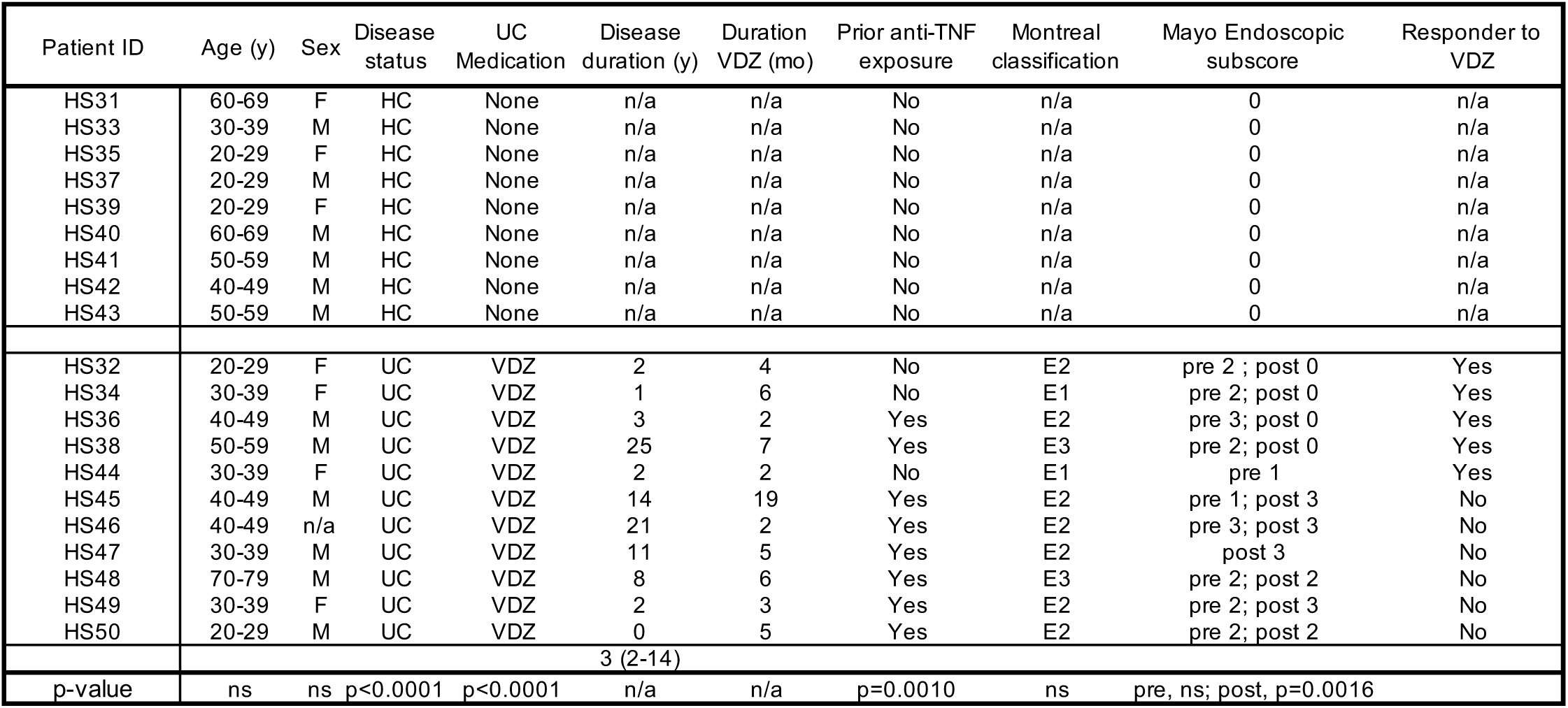
Baseline demographic and clinical data for study participants. Categorical variables were analyzed by Chi-square test and continuous variables were compared using one-way ANOVA with FDR correction or Mann-Whitney test where appropriate. ns, not significant; n/a, not applicable; pre, pre-VDZ treatment; post, post-VDZ treatment. VDZ-vedolizumab.

**Supplementary Table 2.**
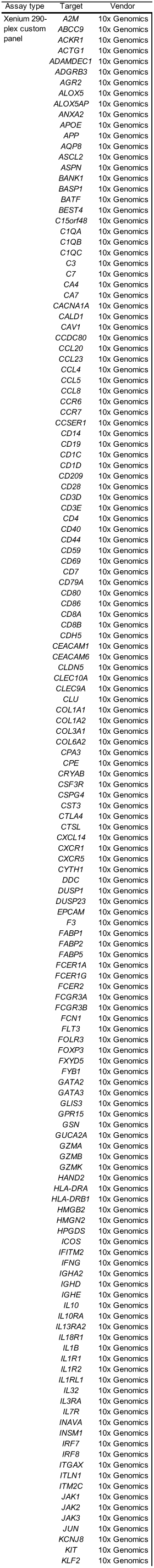
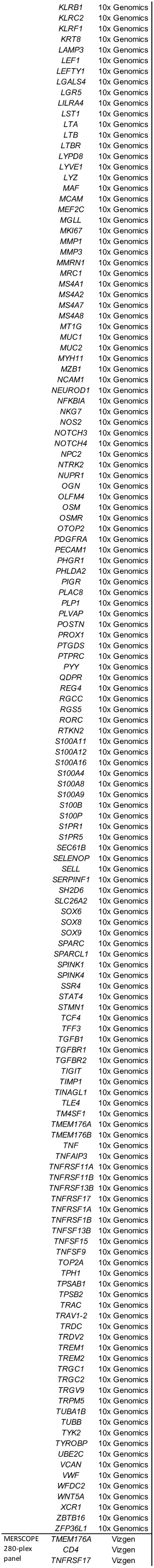
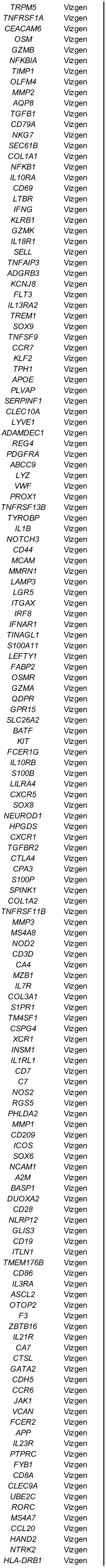
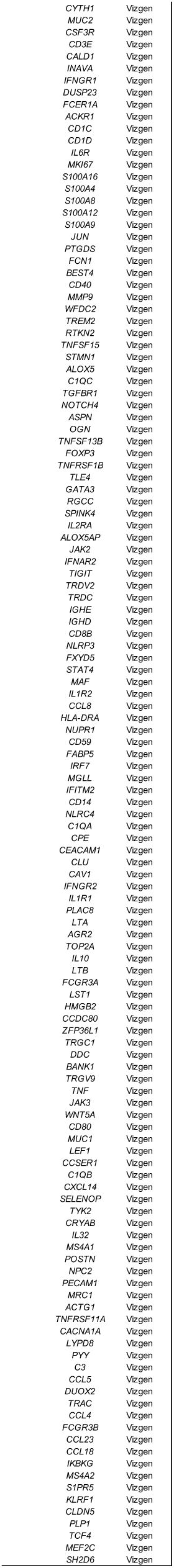
Gene panels for the different spatial transcriptomic platforms.

**Supplementary Table 3.**
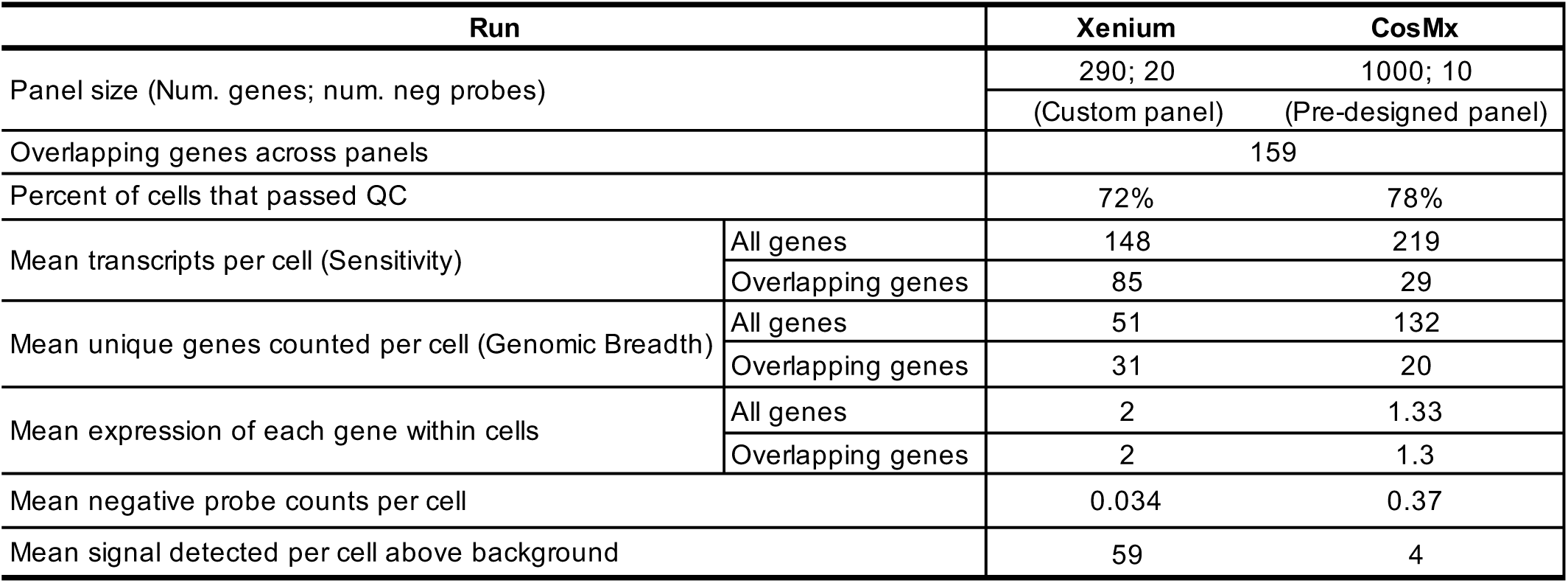
Summarized QC results.

| Run |  | Xenium | CosMx |
| --- | --- | --- | --- |
| Panel size (Num. genes; num. neg probes) |  | 290; 20 | 1000; 10 |
|  |  | (Custom panel) | (Pre-designed panel) |
| Overlapping genes across panels |  | 159 |  |
| Percent of cells that passed QC |  | 72% | 78% |
| Mean transcripts per cell (Sensitivity) | All genes | 148 | 219 |
|  | Overlapping genes | 85 | 29 |
| Mean unique genes counted per cell (Genomic Breadth) | All genes | 51 | 132 |
|  | Overlapping genes | 31 | 20 |
| Mean expression of each gene within cells | All genes | 2 | 1.33 |
|  | Overlapping genes | 2 | 1.3 |
| Mean negative probe counts per cell |  | 0.034 | 0.37 |
| Mean signal detected per cell above background |  | 59 | 4 |

**Supplementary Table 4.**
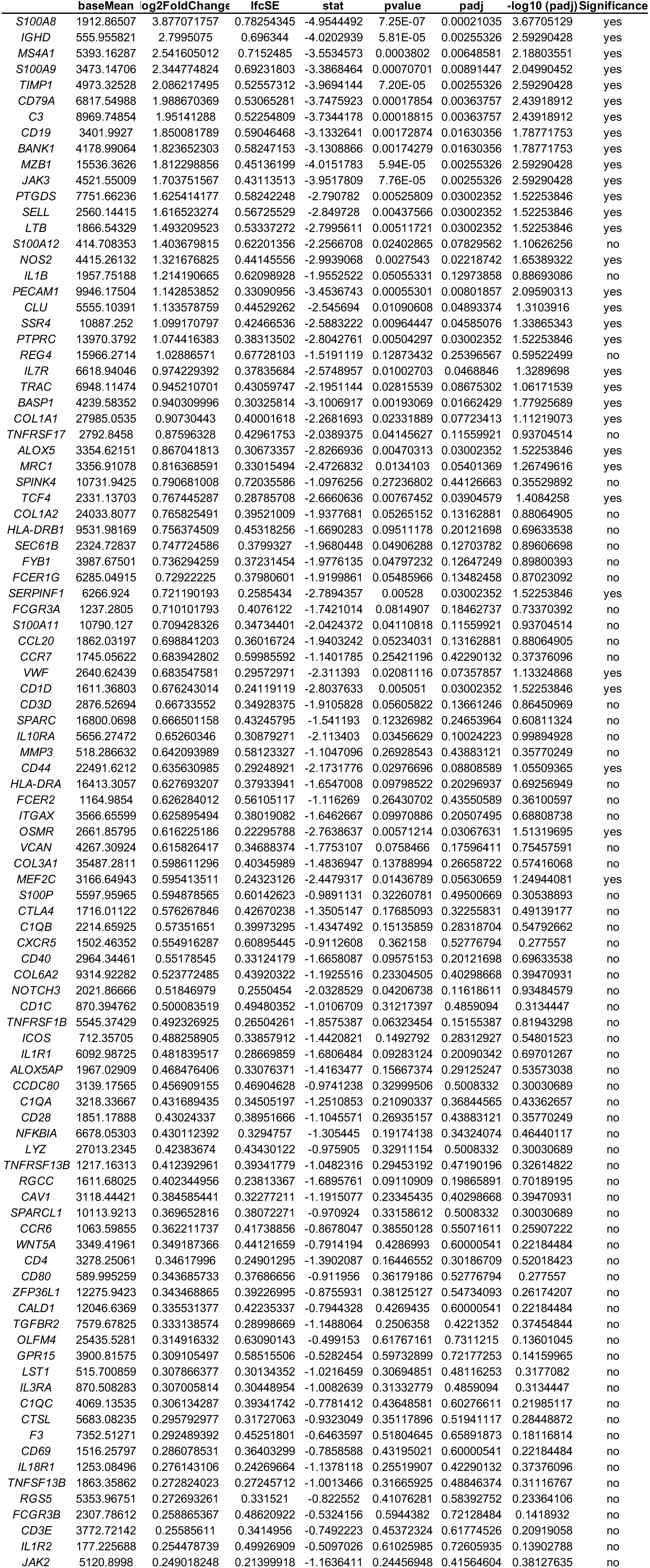
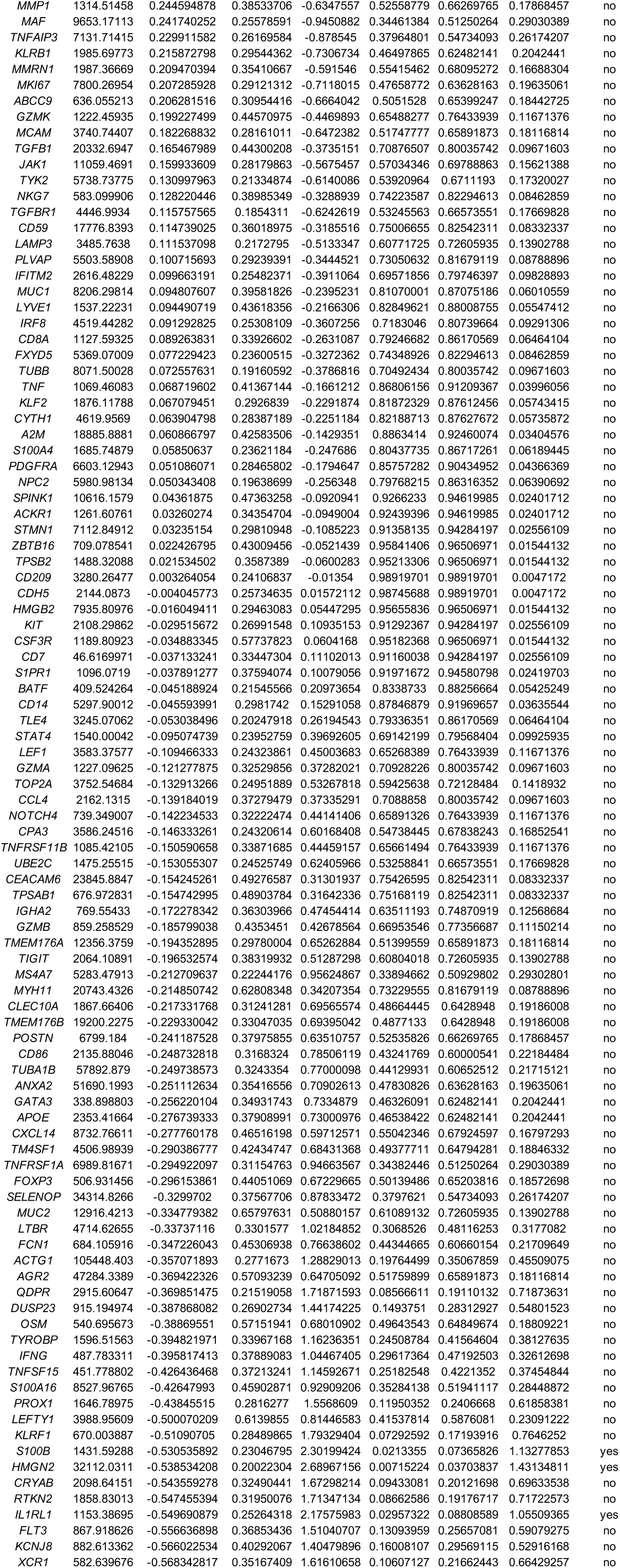
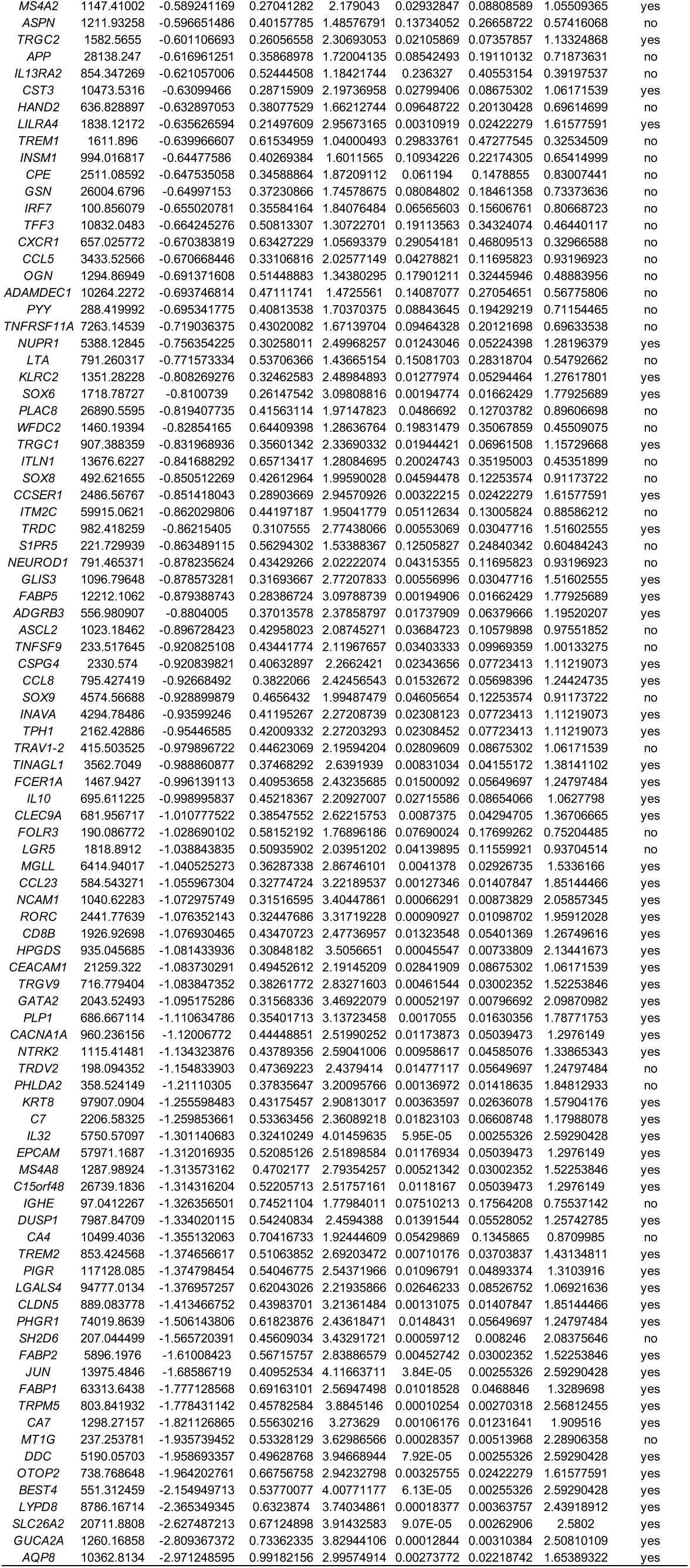
Pseudobulk DE gene analysis of Xenium data comparing colon biopsies in UC PRE versus HC. Significance was set as log2fc >0.4 or <-0.4, p-adj < 0.1 and baseMean >500.

**Supplementary Table 5.**
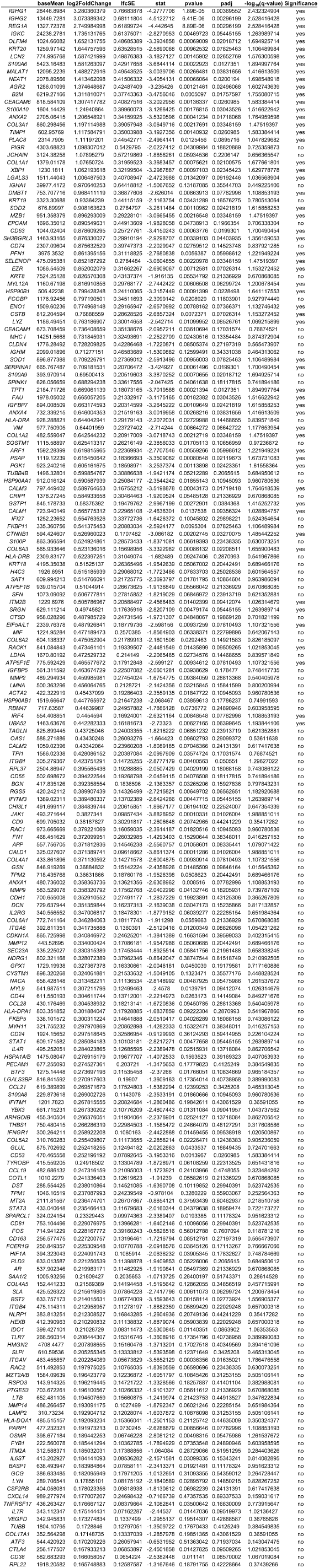
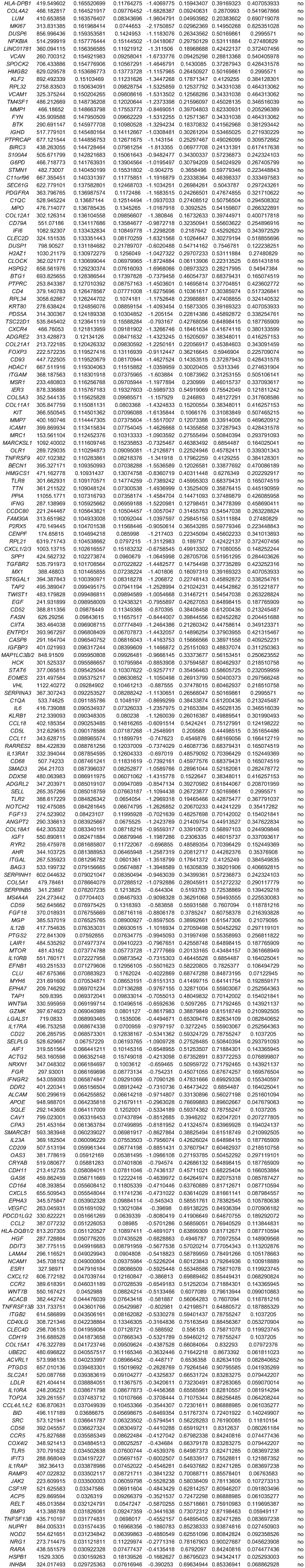
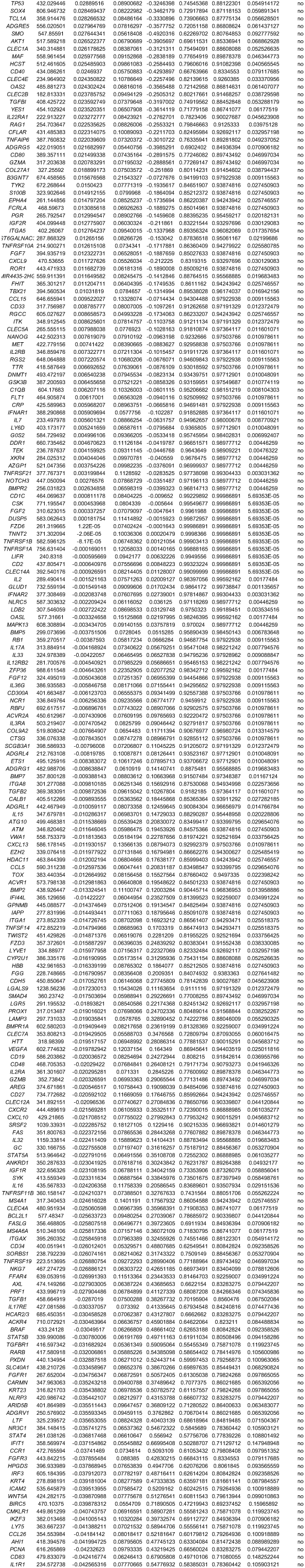
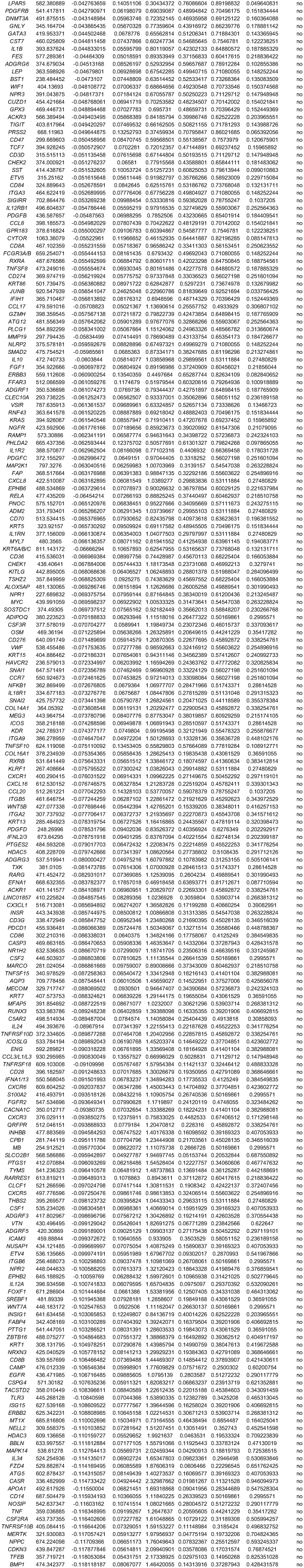
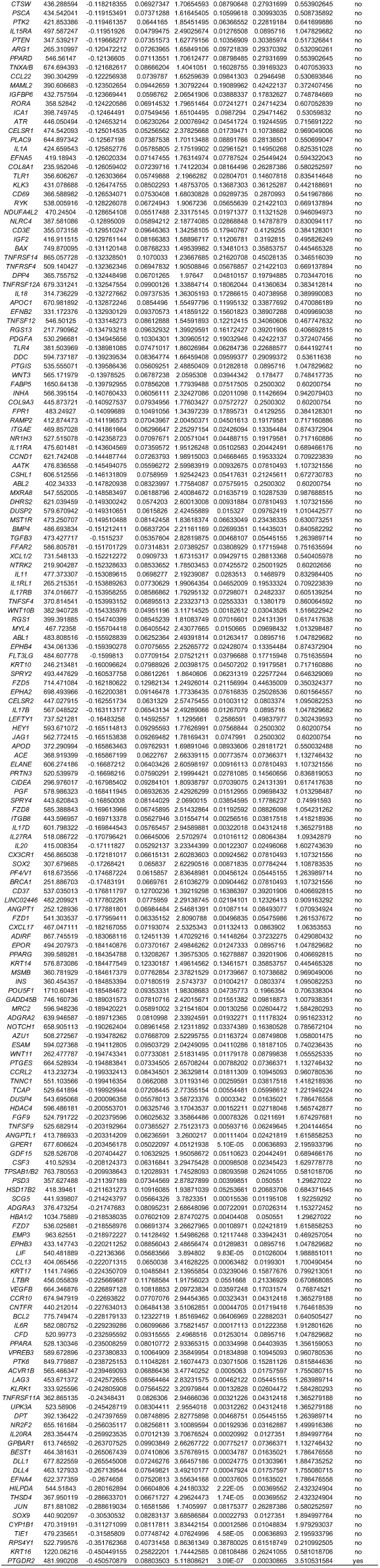
Pseudobulk DE gene analysis of CosMx data comparing colon biopsies in UC PRE versus HC. Significance was set as log2fc >0.4 or <-0.4, p-adj < 0.1 and baseMean >400.

**Supplementary Table 6.**
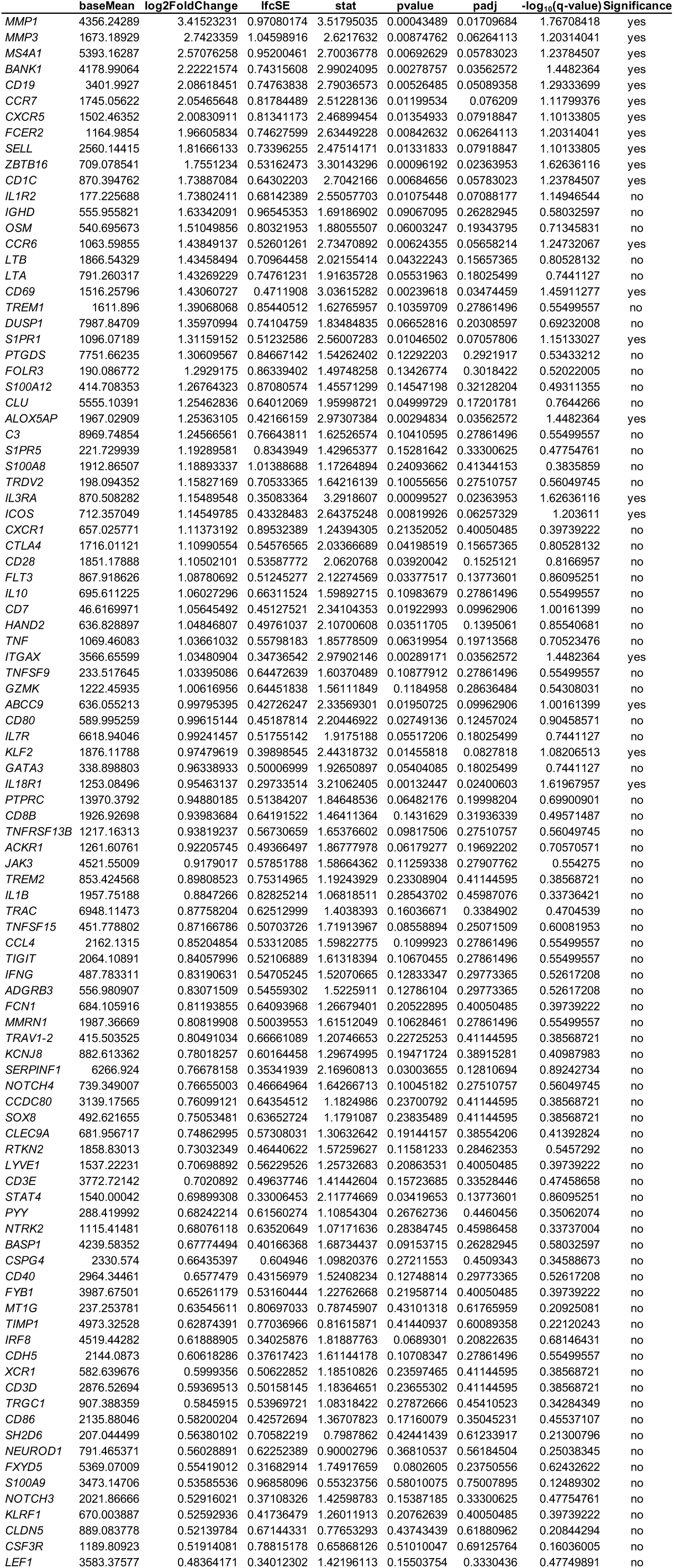
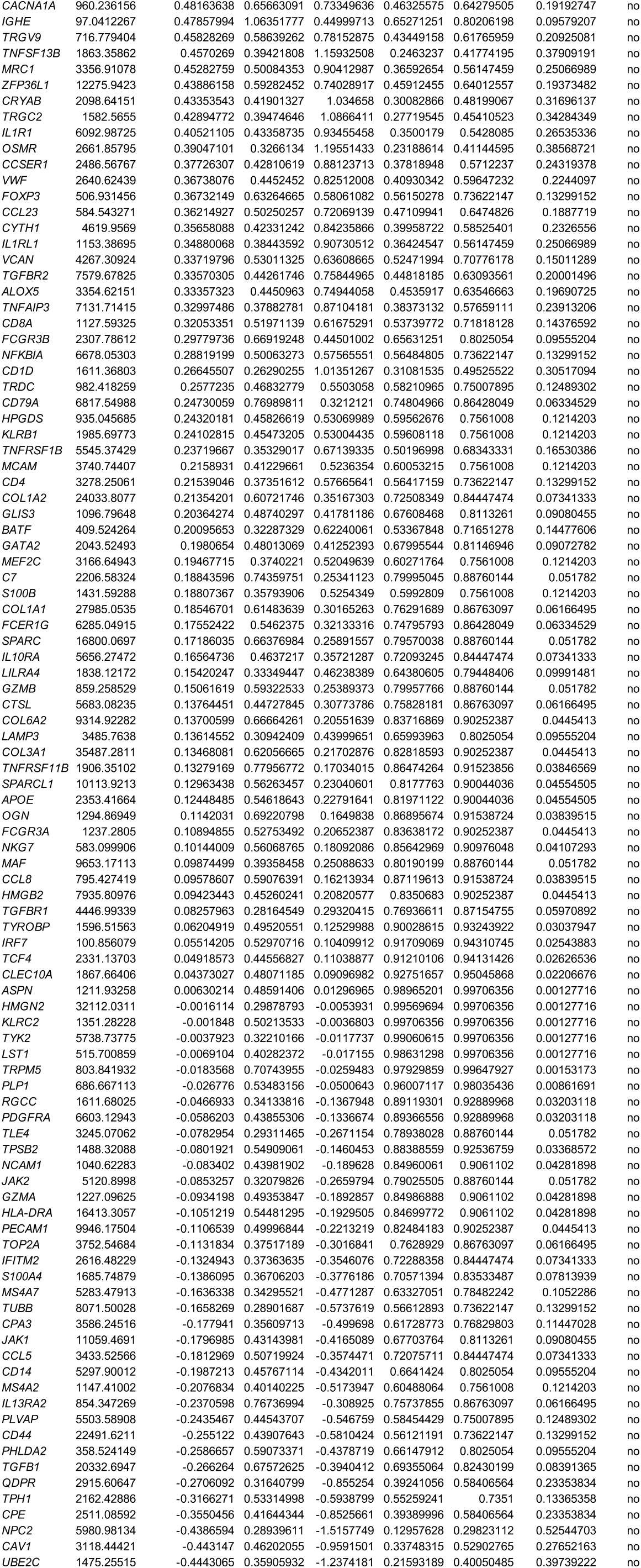
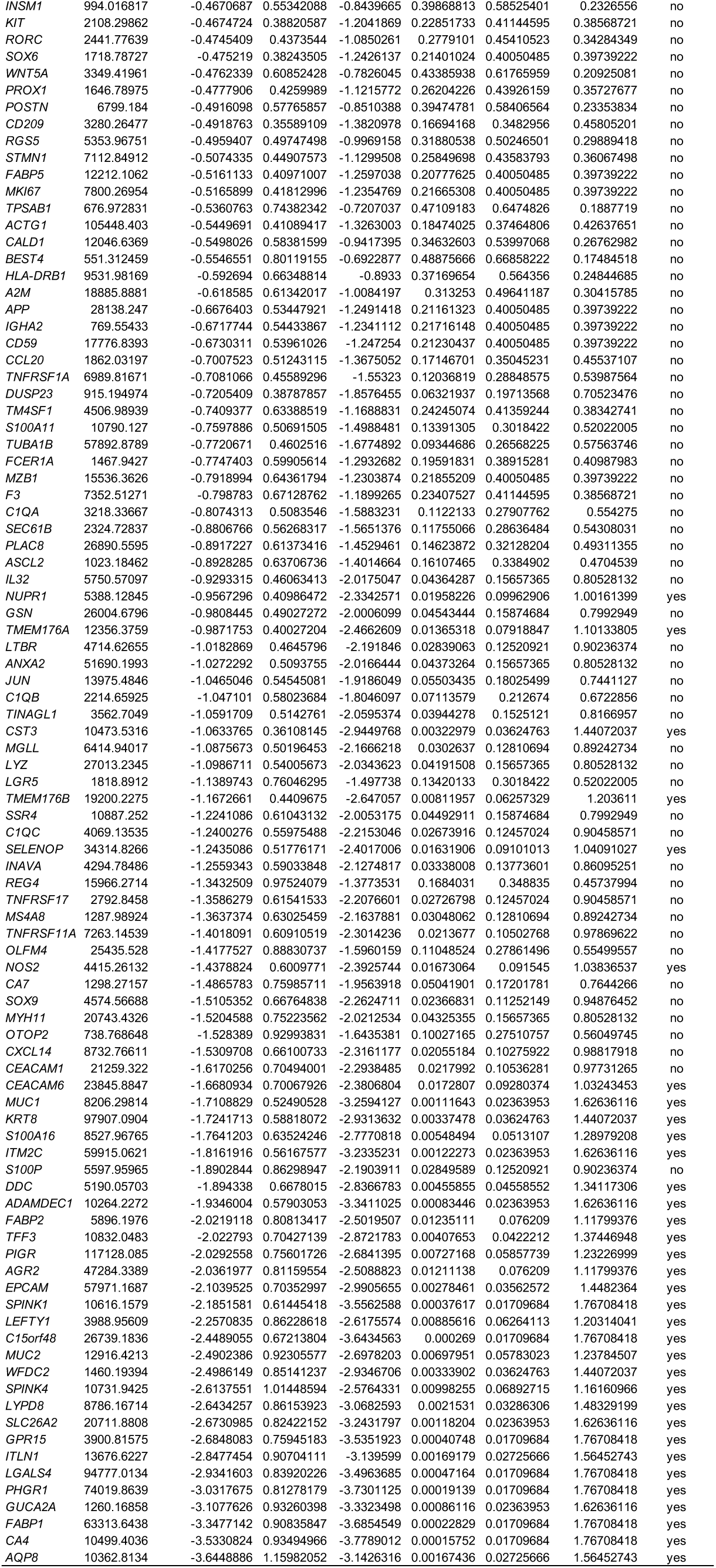
Pseudobulk DE gene analysis of Xenium data comparing colon biopsies in UC PRE Non-Responders versus UC PRE Responders. Significance was set as log2fc >0.4 or <-0.4, p-adj < 0.1 and baseMean >500.

**Supplementary Table 7.**
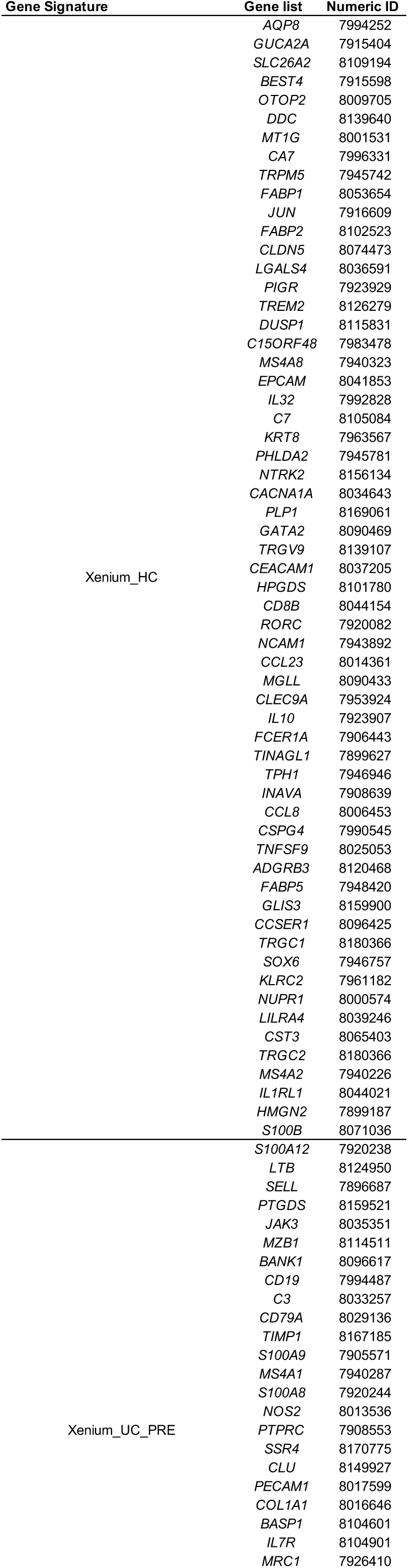
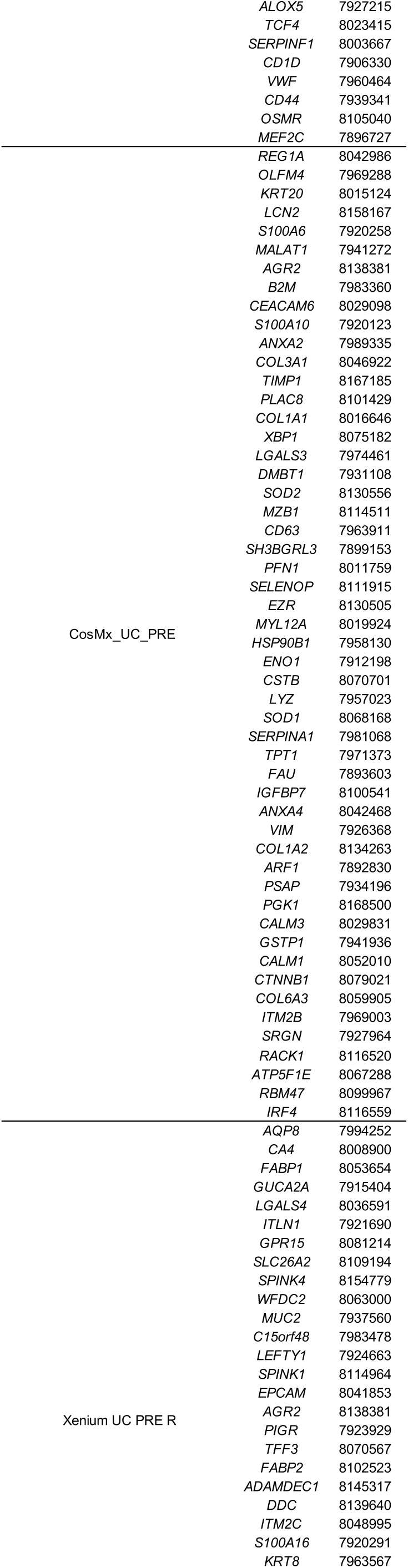
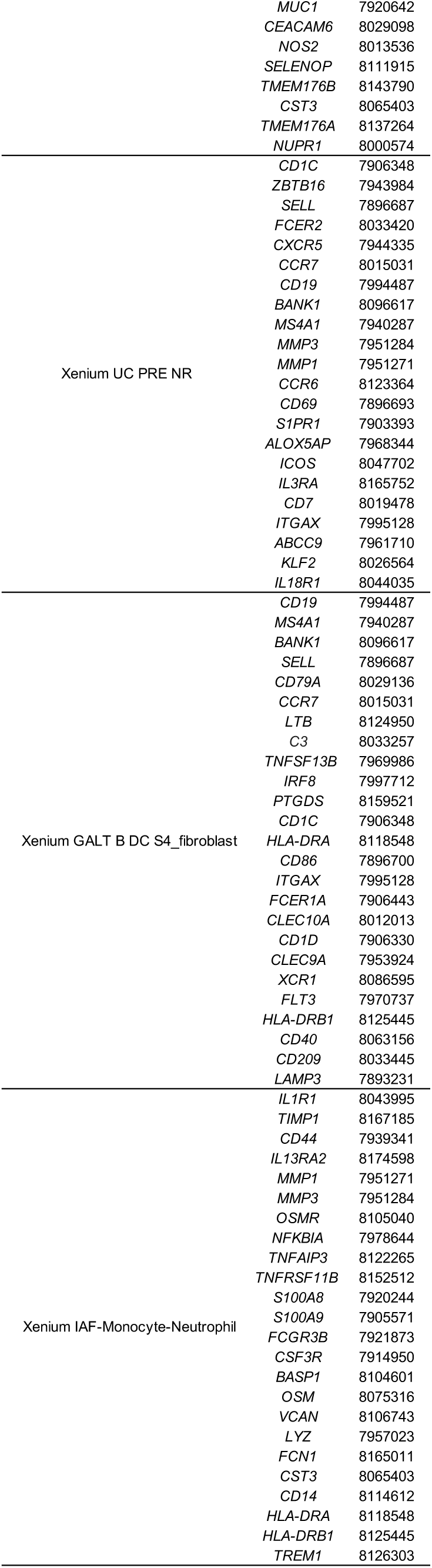
iSCST gene signatures used for Gene Set Enrichment Analysis (GSEA). Numeric ID, Affymetrix numeric probe identifier corresponding to each gene.

